# A Unified Form of Batch Harmonization Equation for Normative Modeling: A Location–Scale Framework

**DOI:** 10.64898/2026.05.17.725713

**Authors:** Min Li, Ying Wang, Yajun Shen, Lin An, Gangyong Jia, Maria L. Bringas-Vega, Pedro Antonio Valdés-Sosa

## Abstract

Normative modeling quantifies individual deviation from population norms by estimating the conditional mean and variance of brain-derived measures as functions of clinically relevant parameters such as age. The rapid growth of multi-center consortia has created an urgent need for normative models that incorporate batch harmonization. Several harmonization methods based on linear mixed models—ComBat, GAMLSS, HBR, and Generalized Normative Modeling (GNM)—offer explicit formulations of the mean and variance, making them natural candidates for batch-harmonized normative modeling; yet the absence of a unified theoretical framework leaves it unclear whether and how these methods support the computation of batch-harmonized *z*-scores. We bridge this gap by writing existing harmonization methods as special cases of a single location–scale equation, *y* = *m*(**x, Θ**)+*σ*(**x, Θ**) *ε*, which we term the *unified form of batch harmonization equation for normative modeling*. The methods differ only in the functional forms of *m* and *σ*, how batch parameters enter **Θ**, and how **Θ** is estimated. This unified form yields both harmonized data *y*\* and site-invariant *z*-scores from the same model, providing a common theoretical language for harmonized normative modeling. Building on this framework, we evaluate the underlying regression engines (parametric, spline, Gaussian process, kernel, deep learning), sensitivity to outliers, computational scalability, and federated decomposability for privacy-preserving multi-center computation. By clarifying what each method assumes, what it delivers, and where the boundaries of current methodology lie, the unified equation establishes a principled foundation for method selection and charts a path toward reliable, scalable, and privacy-aware normative modeling across multi-center neuroimaging.

## 1. Introduction

### 1.1. Normative modeling: from growth charts to brain charts

In clinical medicine, normative modeling has long served as the foundation for individual-level assessment. Pediatric growth charts—relating height, weight, and head circumference to age—have been used for over a century to identify children who deviate from expected developmental trajectories (Cole, 2012). The underlying logic is straightforward: estimate the population distribution of a measure as a function of relevant covariates, then express each individual’s observation as a standardized score (*z*-score) that quantifies their deviation from the norm.

This logic has been extended to neuroscience with considerable success. By constructing normative models of brain-derived descriptive parameters (DPs)— such as cortical thickness, regional volume, or spectral power—as functions of clinically relevant parameters (RCPs) such as age, researchers can characterize individual neurobiological “fingerprints” that go beyond group-level comparisons (Marquand et al., 2016, 2019; Fraza et al., 2021; Bayer et al., 2022). The *z*-score provides individual-level statistical inference: it describes the biological deviation by comparison with the expected reference constructed from population metrics. This approach has been applied to construct developmental brain charts (Bethlehem et al., 2022), detect white matter injury in infants, characterize heterogeneity in schizophrenia and bipolar disorder (Wolfers et al., 2018; Zabihi et al., 2019), build brain-predicted age as a biomarker for neurodegeneration (Cole et al., 2017; Cole and Franke, 2017), and map lifespan trajectories of aperiodic and periodic EEG components (Li et al., 2025).

### 1.2. The multi-site harmonization challenge

Because no single imaging site can provide the sample sizes and demographic diversity needed for accurate normative models, multi-site data pooling has become a necessity. Consortia such as ABIDE (Di Martino et al., 2014), ENIGMA (Thompson et al., 2014), the UK Biobank (Miller et al., 2016), and HarMNqEEG (Li et al., 2022) aggregate thousands of recordings from dozens of sites. However, pooling introduces non-biological sources of variability—differences in recording equipment, acquisition protocols, and preprocessing pipelines—collectively termed “batch effects.” These effects can account for 20–60% of the variance in derived brain measures (Fortin et al., 2018; Pomponio et al., 2020), often exceeding the biological signals of interest and undermining the reproducibility and reliability of results.

In recent years, several harmonization methods have been proposed, broadly falling into three families: extensions of the generalized additive mixed-effect model (GAMM), such as ComBat (Johnson et al., 2007) and GAMLSS (Rigby and Stasinopoulos, 2005); hierarchical Bayesian approaches (Kia et al., 2020, 2022); Generalized Normative Modeling (GNM), which integrates harmonization and normative inference in a single kernel-based location–scale framework (Li et al., 2026a); and machine learning and deep learning methods (Dinsdale et al., 2021; Hu et al., 2024; Lawry Aguila et al., 2022). Among these, methods built on linear mixed models (LMM) and their extensions—ComBat, GAMLSS, and HBR—are a natural fit for batch-effect harmonization in normative modeling, because they directly model the location and scale of the response distribution, which are the same two quantities required to compute a *z*-score.

### 1.3. The disconnect between harmonization and normative modeling

Despite their shared mathematical lineage, LMM-based harmonization methods and normative modeling are usually presented as separate problems with separate solutions. ComBat is described as a “batch effect removal” algorithm that produces harmonized data *y*\*; GAMLSS is described as a flexible regression framework for location, scale, and shape; HBR is described as a Bayesian framework for individualized inference. Each method is published in its own notation, with its own design matrices, link functions, priors, and estimation strategies. As a result, three interrelated problems persist. First, *notational fragmentation*: it is hard to tell, by inspection, which method captures which kind of batch effect, or which assumptions limit each approach, because reviewers and applied researchers must translate between formalisms before they can compare methods on equal footing. Second, the *harmonized-data trap*: ComBat-style methods deliver *y*\* but not *z*-scores, while GAMLSS and HBR can deliver *z*-scores but the literature rarely makes the harmonized-data formula explicit; the relationship between *y*\* and *z*—which model produces site-invariant *z*-scores, which only produces site-adjusted means, and which requires a second normative-modeling step on top of harmonization—is consistently obscured. Third, there is *no common form for the batch-harmonized z-score*: the whole point of running these methods on multi-site data is to obtain a *z*-score that is both calibrated and site-invariant, yet each method derives this score in its own way, making direct comparison impossible. fig. 1 illustrates this disconnect.

**Figure 1:**
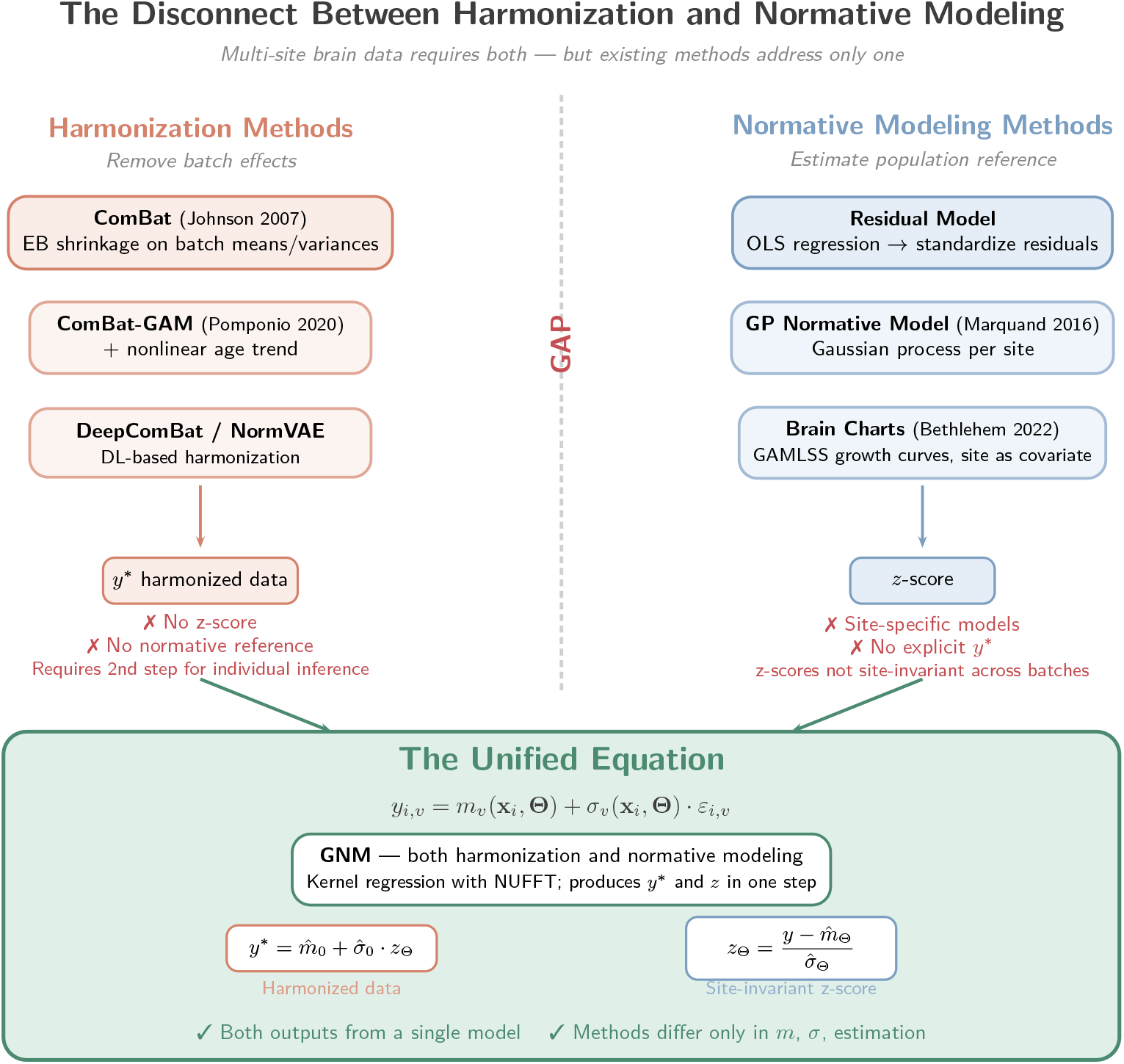
The disconnect between harmonization and normative modeling. Left: harmonization-only methods (e.g., ComBat) produce batch-corrected data *y*\* but no *z*-scores. Right: normative-only methods produce *z*-scores but lack explicit harmonization. Bottom: the unified equation bridges both sides, with GNM as a representative method that simultaneously performs harmonization and normative modeling, yielding both *y*\* and site-invariant *z* from a single location–scale model.

### 1.4. Aim of this review: a unified equation for batch-harmonized z-scores

The aim of this review is to close this gap by writing all major LMM-based harmonization methods in a single, unified normative-modeling form. We show that ComBat, GAMLSS, HBR, and GNM can all be expressed as a single location–scale equation:

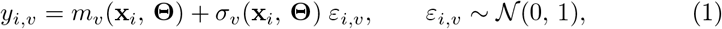

with a single corresponding batch-harmonized *z*-score:

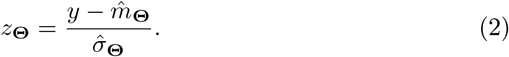

The methods differ only in the functional form chosen for *m*_*v*_ and *σ*_*v*_, in how the batch parameters enter **Θ**, and in how **Θ** is estimated. Once each method is rewritten in this form, both the batch-harmonized *z*-score and the harmonized data *y*\* follow directly and can be compared term-by-term across methods.

This unified form is the central organizing device of everything that follows.We show that all major LMM-based methods are special cases of this equation (section 2), compare the regression engines used to estimate *m*_*v*_ and *σ*_*v*_ (section 3), analyze sensitivity to outliers (section 4), compare computational costs and scalability (section 5), and discuss federated estimation (section 6). We then review validation metrics (section 7) and identify open challenges including multivariate norms, non-Euclidean descriptive parameters, and causal harmonization (section 8).

## 2. The Unified Equation

### 2.1. Notation

Let **Y** *∈* ℝ ^*n×m*^ denote the observation matrix, where *n* is the number of subjects and *m* the number of features (e.g., brain regions, frequency bins). For subject *i* = 1, …, *n* and feature *v* = 1, …, *m*, the observation is *y*_*i,v*_. The corresponding covariate vector is **x**_*i*_ *∈* ℝ ^*p*^ (e.g., age, sex, or jointly frequency and age). The batch (site) identity of subject *i* is denoted *b*_*i*_ *∈ {*1, …, *B}*. table 1 summarizes the notation used throughout this paper.

**Table 1:** Summary of notation.

| Symbol | Description |
| --- | --- |
| $n$ | Number of subjects |
| $m$ | Number of features (brain regions, frequency bins, etc.) |
| $v$ | Feature index, $v = 1, \dots, m$ |
| $y_{i,v}$ | Observation for subject $i$ at feature $v$ |
| $\mathbf{x}_i \in \mathbb{R}^p$ | Covariate vector (e.g., age, sex) |
| $b_i \in \{1, \dots, B\}$ | Batch (site) identity of subject $i$ |
| $B$ | Number of batches (sites) |
| $m_v(\mathbf{x}, \Theta)$ | Location function (conditional mean) |
| $\sigma_v(\mathbf{x}, \Theta)$ | Scale function (conditional standard deviation) |
| $\Theta$ | Full parameter set (fixed + batch effects) |
| $z_{\Theta}$ | Batch-harmonized $z$-score |
| $y^*$ | Harmonized (batch-free) data |
| $K$ | Number of spline basis functions |
| $M$ | Kernel grid size per dimension |
| $d$ | Covariate dimensionality |
| $h$ | Bandwidth (kernel regression) |

### 2.2. From raw standardization to the location–scale form

The element of **Y** can be expressed in terms of its tendency and dispersion:

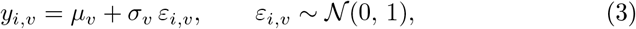

where *µ*_*v*_ and *σ*_*v*_ are the population mean and standard deviation of feature *v*. The conversion of the raw data into the standard score is:

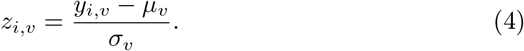

The goal of normative modeling is to quantify individual deviations from the population norm, with accuracy depending on accurate estimates of *µ*_*v*_ and *σ*_*v*_. For Gaussian observations, the *z*-scores are normally distributed, and the normality depends on the estimation accuracy of the mean and standard deviation. The estimation of *µ*_*v*_ = *f*_*v*_(**X**, *θ*) can use smooth algorithms such as splines, local polynomials, or Gaussian processes. The distribution of *σ*_*v*_ is typically heavy-tailed, and a common simplification is to assume homoscedasticity (*σ*_*v*_ = *C*), reducing eq. (3) to a residual model. Marquand et al. (2019) referred to the sample-size-dependent uncertainty as “epistemic” uncertainty 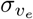and modified the *z*-score as:

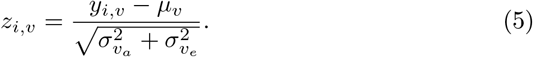

In practice the model is heteroscedastic: the standard deviation depends on covariates. Smooth methods for non-Gaussian distributions or link functions for Gaussianization are needed to estimate *σ*_*v*_ = *g*_*v*_(**X**, *η*). Writing *µ*_*v*_ and *σ*_*v*_ as covariate-dependent functions and adding the batch parameters **Θ** that govern multi-site data, every harmonized normative modeling method can be written as a **location–scale model**:

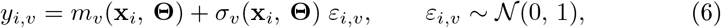

where *m*_*v*_(**x, Θ**) is the **location function** (conditional mean), *σ*_*v*_(**x, Θ**) is the **scale function** (conditional standard deviation), and **Θ** is the full parameter set governing both functions. The corresponding batch-harmonized *z*-score is:

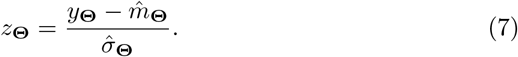

What distinguishes the methods is *how m*_*v*_ and *σ*_*v*_ are parameterized and estimated. In the remainder of this section we show that the residual model, ComBat, HBR, GAMLSS, and GNM are all special cases of eq. (6), and in each case derive both the batch-harmonized *z*-score (eq. (7)) and the harmonized data *y*\* from the same location–scale pair (fig. 2).

**Figure 2:**
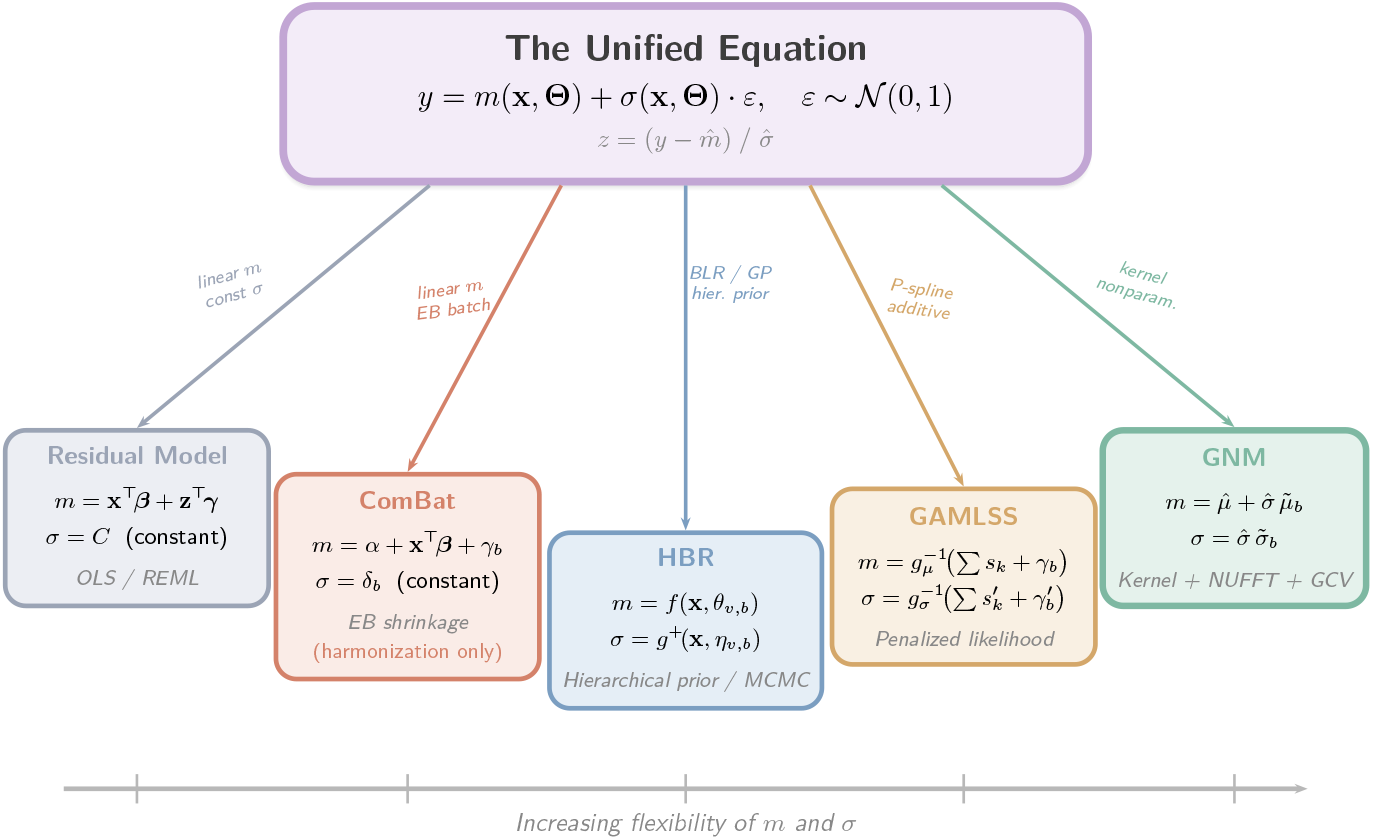
The unified equation as a map of harmonized normative modeling methods. All methods share the same location–scale form (eq. (6)); they differ in the functional flexibility of the location *m*_*v*_ and scale *σ*_*v*_, how batch parameters enter **Θ**, and the estimation strategy.

### 2.3. The residual model

The linear mixed model writes:

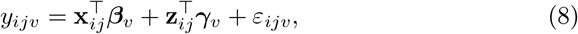

where **x**_*ij*_ is the fixed-effect design vector, **z**_*ij*_ is the random-effect design vector, and ***β***_*v*_ and ***γ***_*v*_ are the corresponding coefficient vectors.

Mapping to eq. (6):

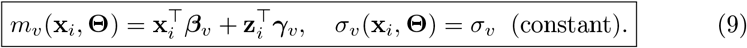

The scale function is **homoscedastic**—it does not depend on **x**.

### 2.4. ComBat: a harmonization-only method

Unlike the other methods, ComBat is *not* a normative modeling method—it is a harmonization method. It removes batch effects from the data but does not directly produce normative *z*-scores. For subject *i* in batch *b* at feature *v*:

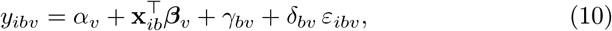

where *α*_*v*_ is the overall mean, ***β***_*v*_ are biological covariate coefficients, *γ*_*bv*_ is the additive batch effect, and *δ*_*bv*_ is the multiplicative batch effect. Johnson et al. (2007) regularize *γ*_*bv*_ and *δ*_*bv*_ via empirical Bayes shrinkage across features.

ComBat cannot directly produce normative *z*-scores for two reasons. Although one could formally write 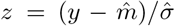, the biological effect **x**^*⊤*^***β***_*v*_ is linear and cannot capture nonlinear normative trajectories; moreover, the scale *δ*_*bv*_ is a batch-specific constant that does not model the population-level heteroscedastic variance *σ*(**x**).

In practice, ComBat is therefore used as a preprocessing step. The batch-adjusted data is:

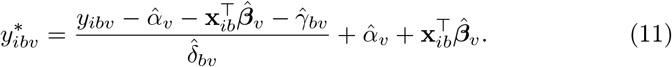

A *separate* normative model is then fitted to *y*\* to estimate *µ*(**x**) and *σ*(**x**), and the *z*-score is computed as 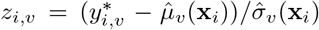. This two-step nature means that harmonization quality and normative estimation accuracy are decoupled—errors in one step propagate to the next but cannot be jointly optimized.

### 2.5. Hierarchical Bayesian Regression (HBR)

Kia et al. (2020, 2022) proposed a hierarchical Bayesian framework for multi-site normative modeling. The mean and scale are estimated as functions *f*_*v*_(**x**, *θ*) and 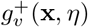, with hierarchical priors on the batch-specific parameters:

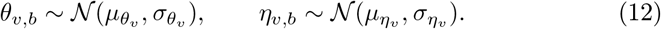

Each batch has its own parameters *θ*_*v,b*_ and *η*_*v,b*_, drawn from a common population distribution, enabling information sharing across batches.

Mapping to eq. (6):

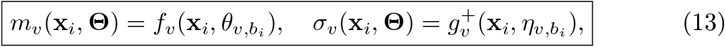

where 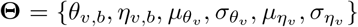. The functions *f*_*v*_ and 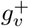can be non-linear (Bayesian linear regression with basis functions, Gaussian processes). The hierarchical prior provides regularization: batch-specific parameters are shrunk toward the population mean, which is particularly beneficial for small-batch samples.

The batch-harmonized *z*-score uses batch-specific parameters, absorbing the site effect:

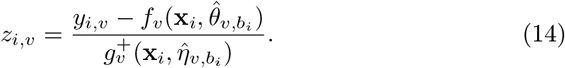

Harmonized data are reconstructed by projecting through the population-level parameters:

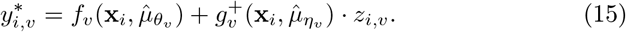

### 2.6. GAMLSS

Rigby and Stasinopoulos (2005) developed the generalized additive model for location, scale, and shape (GAMLSS). Each distribution parameter *k* = 1, …, *K* is modeled through a link function *g*_*k*_(·):

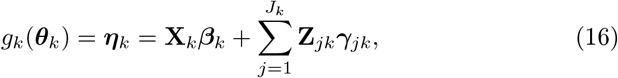

where **X**_*k*_ is the fixed-effect design matrix, ***β***_*k*_ the coefficient vector, **Z**_*jk*_ the random-effect design matrix, and ***γ***_*jk*_ the random-effect vector. For a four-parameter distribution (location *µ*, scale *σ*, skewness *ν*, kurtosis *τ*), each parameter has its own linear predictor. The site enters as a random effect in the linear predictor for each parameter.

Mapping to eq. (6), focusing on the location and scale parameters needed for normative modeling:

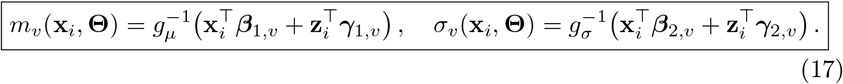

The design matrices can include smooth terms (P-splines) for nonlinear covariate effects.

The batch-harmonized *z*-score is:

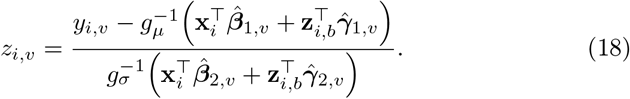

Harmonized data are obtained by projecting the *z*-score through fixed effects only (without site random effects):

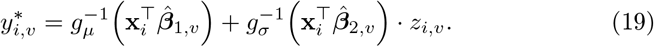

### 2.7. GNM

GNM uses a two-level hierarchical structure. The observation is:

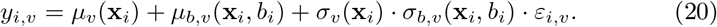

To facilitate hierarchical *z*-score computation, the batch effects are rescaled by the global standard deviation. Defining 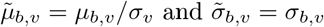:

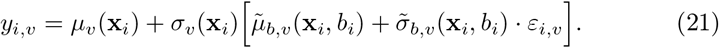

The mean and log-scale functions decompose additively:

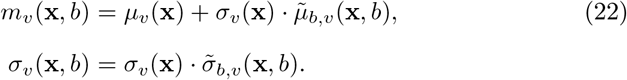

This equation encompasses ComBat and GAMLSS as special cases: when the fixed-effect parameters are linear and the random-effect parameters are constant, it reduces to ComBat; when the random-effect parameters are linear in covariates, it reduces to GAMLSS.

Mapping to eq. (6):

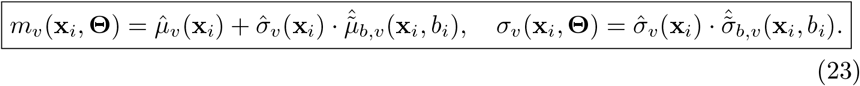

All functions are estimated nonparametrically via kernel regression with NUFFT acceleration and GCV bandwidth selection.

GNM computes the *z*-score hierarchically:

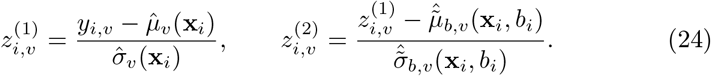

Harmonized data are obtained by projecting the final *z*-score through only the global-level parameters:

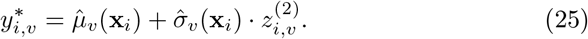

### 2.8. Summary comparison

table 2 presents a term-by-term comparison of the five methods under the unified equation. Each row shows how a method instantiates the location function *m*_*v*_, scale function *σ*_*v*_, batch-effect handling, estimation strategy, and whether it directly produces a *z*-score. fig. 3 visualizes the subsumption relationships: the residual model is nested within ComBat, which is nested within GAMLSS, which is nested within GNM; HBR forms a parallel Bayesian branch.

**Table 2:**
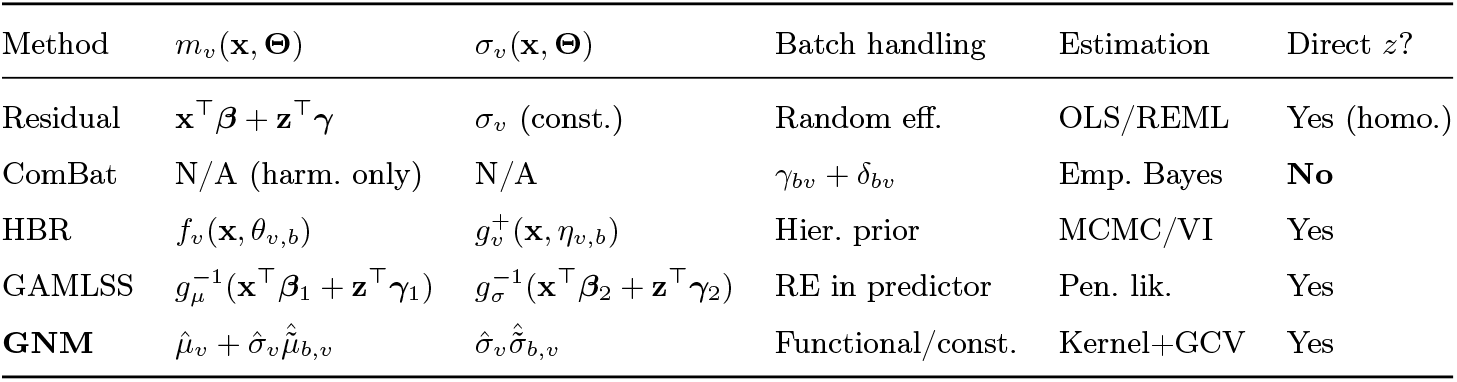
Summary comparison of harmonized normative modeling methods under the unified equation (eq. (6)).

**Figure 3:**
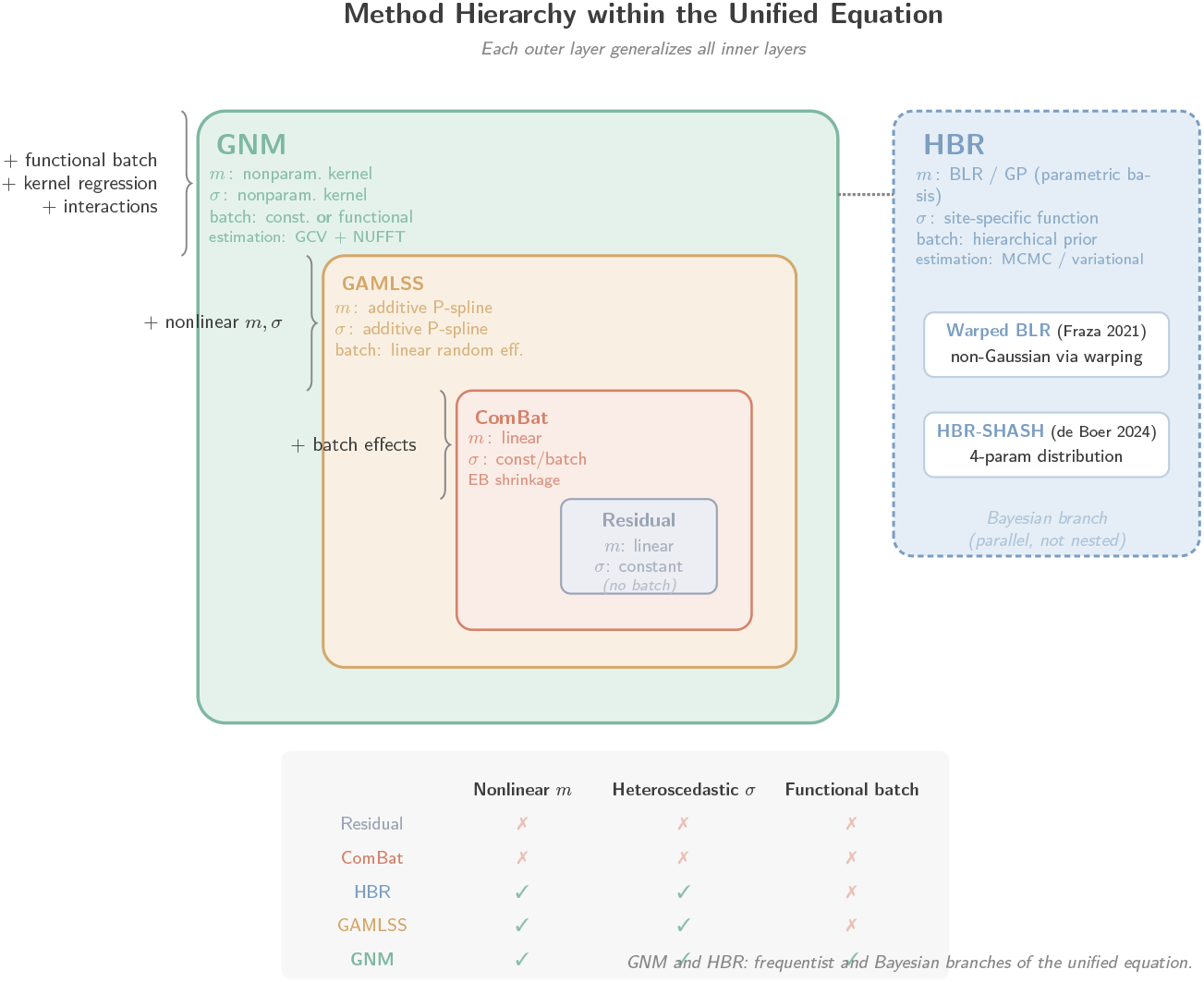
Method hierarchy within the unified equation. Each outer layer generalizes all inner layers: the residual model (homoscedastic, no batch) is nested within ComBat (adds batch effects), which is nested within GAMLSS (adds nonlinear location and scale), which is nested within GNM (adds functional batch effects and kernel regression). HBR forms a parallel Bayesian branch. GNM and HBR represent the frequentist and Bayesian branches of the unified equation.

## 3. Regression Methods Underlying the Unified Equation

The unified equation (eq. (6)) specifies the *form* of the normative model but not *how* the location and scale functions are estimated. The choice of regression engine determines the model’s flexibility, computational cost, and sensitivity to model misspecification. We review the major families of estimators used across the methods discussed in section 2; a more detailed treatment of regression algorithms for normative modeling is given in Li et al. (2026b).

### 3.1. Parametric methods: OLS and polynomial regression

The simplest approach models *m*_*v*_(**x**) as a linear or polynomial function of covariates: *m*_*v*_(**x**) = **x**^*⊤*^***β***_*v*_. This is the backbone of the residual model (eq. (8)) and ComBat (eq. (10)). Estimation is by ordinary least squares (OLS) or restricted maximum likelihood (REML) for mixed models, with computational cost *O*(*np*^2^) where *p* is the number of regressors. Parametric models are fast, interpretable, and statistically efficient when the assumed form is correct; empirical Bayes shrinkage, as in ComBat, further stabilizes estimates for small batches by borrowing strength across features. These properties made the parametric family the default choice in the first generation of multi-site harmonization.

The limitation is that linear models cannot capture the nonlinear normative trajectories that are ubiquitous in neuroscience—for example, the inverted-U shape of cortical thickness across the lifespan (Bethlehem et al., 2022). Polynomial extensions such as quadratic or cubic regression offer limited flexibility and suffer from Runge’s phenomenon at the boundaries of the covariate range, where neuroimaging data are also sparsest. The scale function *σ*_*v*_ is typically assumed constant (homoscedastic), which biases *z*-scores when inter-individual variability changes with age. Under this framework, *m*_*v*_ is a linear function of **x** and *σ*_*v*_ is a covariate-independent constant—the most restrictive specialization.

### 3.2. Spline-based methods: P-splines, B-splines, and GAM

Spline regression represents *m*_*v*_(**x**) as a linear combination of basis functions (B-splines or P-splines) placed at fixed knots along the covariate axis. Penalized splines (P-splines; Eilers and Marx, 1996) add a roughness penalty to prevent overfitting, with the penalty weight selected by cross-validation or REML. GAMLSS (section 2.6) uses P-splines as the default smooth term, modeling each distribution parameter as an additive function *g*_*k*_(*θ*_*k*_) = Σ_*j*_ *s*_*j*_(*x*_*j*_) + random effects, so that both the location and scale functions become covariate-dependent. The Brain Charts project (Bethlehem et al., 2022) applied GAMLSS with P-spline smoothers at unprecedented scale (~120,000 MRI scans), demonstrating that spline methods are both flexible enough for lifespan neurodevelopment and scalable enough for consortium-size data.

The cost of this flexibility is primarily structural. The additive form Σ_*j*_ *s*_*j*_(*x*_*j*_) does not capture interactions between covariates unless explicit tensor product terms are added, and the size of the tensor-product basis grows combinatorially with dimension. Knot placement, while largely automated by P-spline penalties, still requires the user to specify the number and range of basis functions, and default choices may break down when the sampling density is uneven. For multivariate covariates—for example, joint frequency–age surfaces in EEG spectral power—the additive approximation can be inadequate and the analyst is pushed toward less interpretable tensor-product specifications. Computationally, a single backfitting iteration runs in *O*(*NK*^2^) with *K* basis functions, which remains tractable well into the tens of thousands of subjects.

### 3.3. Bayesian nonparametric methods: Gaussian processes and BLR

Hierarchical Bayesian regression (section 2.5) can use either Bayesian linear regression (BLR) with fixed basis functions or Gaussian process (GP) regression. In BLR,

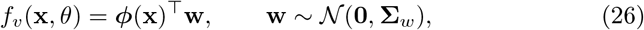

where ***ϕ***(**x**) is a basis expansion (polynomial, B-spline). In GP regression,

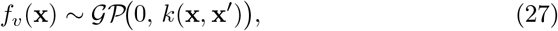

with a kernel function *k* such as the squared exponential. Fraza et al. (2021) extended BLR with a warping function to handle non-Gaussian responses (Warped BLR), while de Boer et al. (2024) introduced the sinh-arcsinh (SHASH) likelihood within HBR, modeling all four distribution parameters—location, scale, skewness, and kurtosis—as functions of covariates and site. The Bayesian framework gives principled uncertainty quantification because the posterior predictive distribution yields both point estimates and credible intervals, and the hierarchical prior on batch parameters provides automatic shrinkage for small sites. GP regression is fully nonparametric and in principle captures arbitrary smooth functions.

These benefits come at a substantial cost. Exact GP regression requires *O*(*N* ^3^) time and *O*(*N* ^2^) memory, making it impractical for large datasets (more than about 10,000 subjects) without sparse approximations based on inducing points, which themselves introduce approximation error. MCMC inference is slow and requires careful convergence diagnostics, and variational approximations trade some of the Bayesian rigor for tractability. BLR with fixed basis functions inherits the limitations of the chosen basis—B-splines cannot capture interactions without tensor products, polynomials suffer from the same boundary problems discussed above—so the analyst again faces the flexibility–dimensionality tradeoff, now wrapped in a Bayesian posterior.

### 3.4. Kernel methods: Nadaraya–Watson, LPR, and NUFFT acceleration

Kernel regression estimates *m*_*v*_(**x**) as a locally weighted average of the data.

The Nadaraya–Watson (NW) estimator is

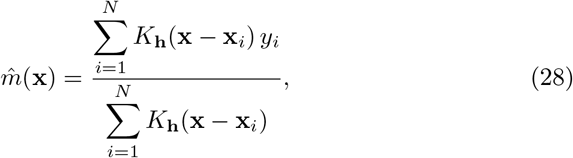

where *K*_**h**_ is a product kernel with bandwidths **h**. Local polynomial regression (LPR; Fan and Gijbels, 1996) generalizes NW by fitting a polynomial of order *l* within each kernel window, reducing boundary bias. GNM (section 2.7) uses LPR with the Epanechnikov kernel, accelerated by the Non-Uniform Fast Fourier Transform (NUFFT). The NUFFT acceleration exploits the fact that the kernel-weighted sums in LPR are discrete convolutions: by projecting non-uniformly sampled data onto a regular grid via fast Gaussian gridding, computing the convolution in the frequency domain, and interpolating back, the cost is reduced from *O*(*N* ^2^) to *O*(*N* + *M* ^*d*^ log *M*), where *M* is the grid size per dimension and *d* is the covariate dimension (Greengard and Lee, 2004; Wang et al., 2022, 2026; Li et al., 2026a). Because the framework treats *m*_*v*_ and *σ*_*v*_ symmetrically, the same engine estimates both functions, and the same NUFFT acceleration applies to the mean and variance smoothers alike. Kernel methods are therefore fully nonparametric, naturally multivariate, capable of capturing interaction effects without explicit specification, and equipped with a single hyperparameter—the bandwidth—that can be selected automatically by generalized cross-validation (GCV).

Two caveats temper this flexibility. Kernel methods suffer from the curse of dimensionality: performance degrades rapidly for *d >* 3, so the framework is best suited to one-to three-dimensional covariate sets such as age, or jointly frequency and age. Bandwidth selection is global—one bandwidth per covariate dimension—which can be suboptimal when the smoothness of the underlying function varies across the covariate space, and boundary effects require deliberate handling (for example, linear extrapolation outside the observed covariate range in GNM). In practice, neuroscience applications of normative modeling rarely need more than a handful of continuous covariates, so these caveats are manageable; when they matter, they motivate extensions such as adaptive-bandwidth kernels rather than a shift to a different estimator family.

### 3.5. Deep learning methods

Deep learning approaches to harmonization and normative modeling attempt to learn flexible, high-dimensional mappings that disentangle biological signal from site effects. DeepComBat (Hu et al., 2024) uses a conditional variational autoencoder to learn site-invariant representations while preserving biological covariates. Adversarial unlearning (Dinsdale et al., 2021) augments a prediction network with an adversary that attempts to recover site identity from learned features, pushing the encoder toward site-invariant representations. Conditional VAEs (Moyer et al., 2020; Lawry Aguila et al., 2022) model the generative process of brain images conditioned on covariates and site, learning to separate conditional sources of variation in the latent space.

The appeal of these methods lies in their capacity to capture nonlinear interactions between covariates and site effects and to operate directly on high-dimensional inputs (voxel-wise data, dense connectivity matrices) without a prior feature extraction step. However, they sit uneasily within the unified equation (eq. (6)) because they do not expose a closed-form location–scale pair. The conditional mean and variance are implicit in a learned latent representation rather than in explicit functions *m*_*v*_(**x, Θ**) and *σ*_*v*_(**x, Θ**), making it difficult to compute calibrated *z*-scores: the latent codes do not, in general, follow a known distribution conditional on covariates, and probing the network at a specific **x** does not cleanly separate the conditional mean from the conditional variance. Training requires large datasets and careful hyperparameter tuning, and reproducibility and uncertainty quantification remain open challenges. Deep learning methods are therefore best viewed as complementary to the unified location–scale framework: they are most useful when the data structure defeats explicit parametric or nonparametric regression, but they sacrifice the interpretability and calibration guarantees that the unified equation provides.

### 3.6. Summary: regression method comparison

table 3 summarizes the key properties of each regression engine.

**Table 3:** Comparison of regression engines used in normative modeling and harmonization.

| Regression engine | Flexibility | Multivariate | Computational cost | Used by |
| --- | --- | --- | --- | --- |
| OLS / polynomial | Low | Yes (linear) | $O(np^2)$ | Residual, ComBat |
| P-spline / GAM | Medium (additive) | Additive only | $O(NK^2)$ | GAMLSS |
| GP | High | Yes | $O(N^3)$ | HBR |
| BLR + basis | Medium | Depends on basis | $O(Np^2)$ | HBR |
| Kernel / LPR + NUFFT | High | Yes (product kernel) | $O(N + M^d \log M)$ | GNM |
| Deep learning | Very high | Yes | $O(N \cdot \text{epochs})$ | DeepComBat, NormVAE, Adversarial |

## 4. Outlier Influence

Outliers are a pervasive concern in multi-site neuroimaging data. They arise from motion artifacts, preprocessing failures, scanner malfunctions, data entry errors, and genuine biological extremes. Because normative modeling estimates both the mean *m*_*v*_(**x**) and the variance *σ*_*v*_(**x**), outliers contaminate both estimates—but at different rates depending on the estimator. The accuracy of outlier handling is particularly important because the *z*-score interpretation relies on a well-calibrated reference distribution: a model contaminated by outliers will systematically misclassify clinical deviations (Marquand et al., 2019, 2014). Outlier proportions in published multi-site neuroimaging studies typically range from 2% to 10% (Wager et al., 2014; Pomponio et al., 2020), well within the range that compromises classical least-squares estimators.

### 4.1. Comparative sensitivity across methods

The five families of estimators discussed in section 2 span a wide range of outlier sensitivity (fig. 4), which can be understood by tracking how a single contaminated observation propagates through the location and scale estimates. At one extreme, OLS-based methods—including the residual model and ComBat—have a breakdown point of 1*/n*, meaning that a single sufficiently extreme observation can shift 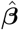arbitrarily (Hampel, 1971). In ComBat, contamination enters at two levels: the global regression for ***β***_*v*_ and the batch-level estimation of *γ*_*bv*_ and *δ*_*bv*_. Although empirical Bayes shrinkage stabilizes the batch parameters by borrowing strength across features (Johnson et al., 2007), it operates on the parameters rather than the data points, so a single extreme observation in a small batch can still dominate the batch mean. Pomponio et al. (2020) reported that ComBat required pre-screening of subjects with extreme *z*-scores in the Lifespan Brain Chart data, and Horng et al. (2022) found that ComBat variants (CovBat, ComBat-GAM) remain vulnerable in high-variance features.

**Figure 4:**
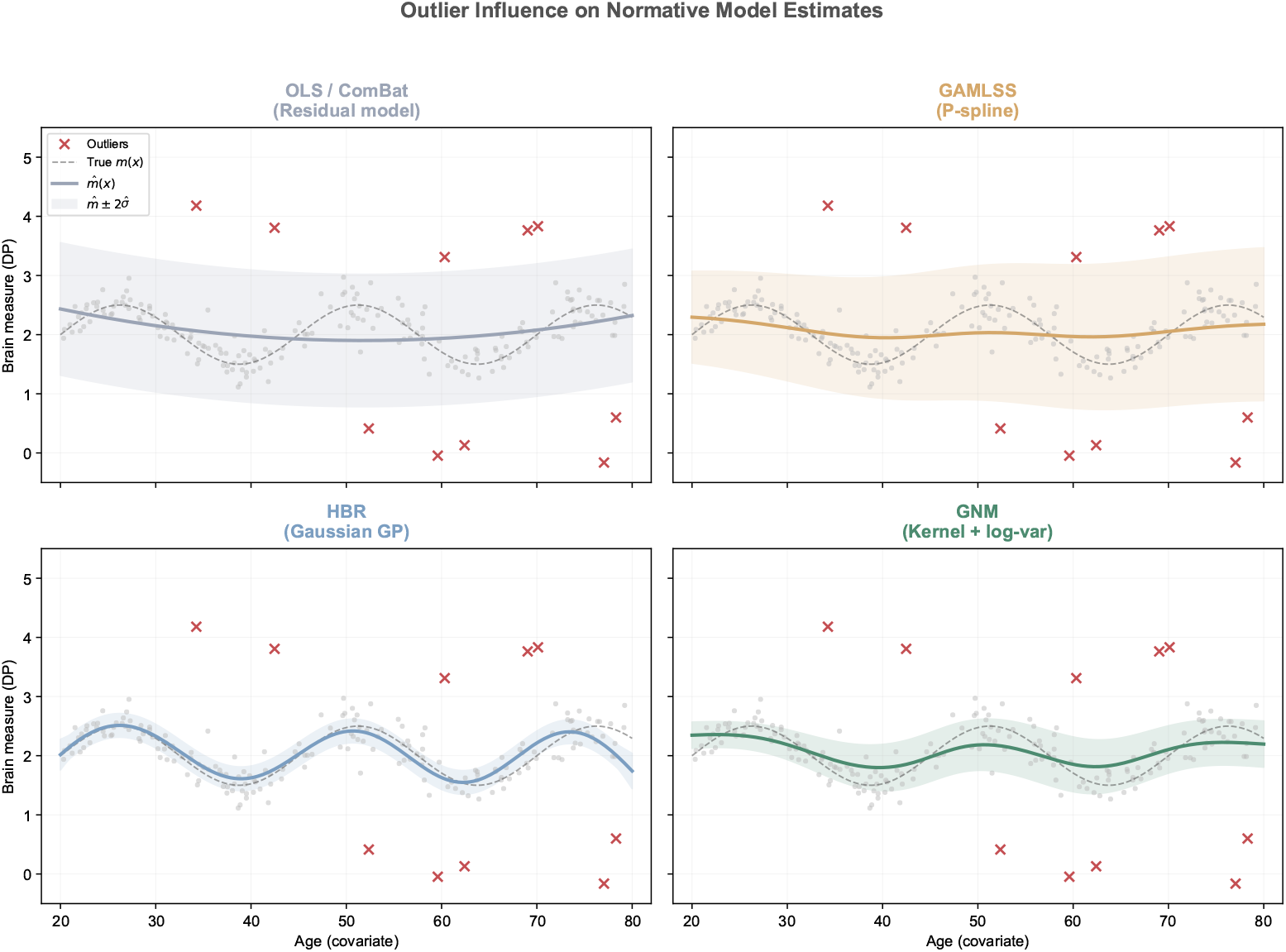
Outlier influence on normative model estimates. Four methods are compared on synthetic data with 5% outlier contamination (red crosses). OLS/ComBat (top left): the quadratic fit is pulled toward outliers, producing a wide, poorly calibrated confidence band. GAMLSS with P-splines (top right): the smooth mean resists isolated outliers, but the variance estimator inflates substantially because squared residuals amplify extreme values. HBR with Gaussian GP (bottom left): the flexible posterior mean tracks outliers locally; the Gaussian likelihood provides no down-weighting mechanism. GNM with kernel regression and log-space variance (bottom right): the local averaging dilutes outlier influence on the mean, and log-space variance estimation compresses extreme positive residuals, yielding the tightest calibrated band.

Penalized likelihood methods such as GAMLSS occupy a middle ground. The P-spline roughness penalty prevents the smooth function from interpolating individual data points, conferring moderate resistance to isolated outliers in the location estimate (Eilers and Marx, 1996). However, the scale estimate 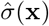 is based on squared residuals, which amplify extreme values and inflate the estimated variance; the result is deflated *z*-scores and reduced sensitivity to genuine deviations (Stasinopoulos and Rigby, 2007). This vulnerability is exacerbated when outliers cluster at particular covariate values—for instance, older subjects with undetected pathology—where both 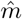and 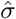can be locally biased. GAMLSS addresses non-Gaussian residuals through its distribution family: the Box-Cox-*t* (BCT) distribution combines a power transformation for skewness with Student-*t* tails for heavy-tailed contamination, and Bethlehem et al. (2022) found that BCT and BCPE distributions outperformed Gaussian fits in brain regions with skewed distributions.

Hierarchical Bayesian regression provides implicit regularization through the hierarchical prior, which shrinks batch-specific parameters toward the population mean and thereby reduces the influence of outliers in small batches (Kia et al., 2020, 2022). Nonetheless, the standard Gaussian likelihood offers no mechanism for down-weighting observations far from the conditional mean. Two extensions address this limitation. The Student-*t* likelihood places a posterior on the degrees of freedom *ν*, effectively learning the contamination level from the data and assigning lower weight to extreme observations (Fernandez and Steel, 1998; Geweke, 1993). The sinh-arcsinh (SHASH) likelihood, introduced by de Boer et al. (2024) within HBR, goes further by separating skewness from tail weight, allowing the model to accommodate asymmetric heavy tails. In their evaluation on an OpenNeuro multi-site dataset, SHASH-HBR produced better-calibrated *z*-scores than Gaussian HBR, particularly in subcortical regions with skewed distributions.

Kernel regression, as used in GNM, derives its resistance from local averaging: the Nadaraya–Watson estimator dilutes the influence of individual outliers across all observations within the bandwidth window, with larger bandwidths providing greater dilution (Fan and Gijbels, 1996). The variance estimator, however, remains sensitive because squaring amplifies extreme values. GNM partially mitigates this through log-space variance estimation (Chen and Bickel, 2006), which compresses extreme positive residuals. Local M-regression—replacing the local least-squares objective with a bounded loss such as Huber or Tukey bisquare—has been studied in the statistical literature (Härdle, 1990; Fan, 1993) but has not yet been integrated into neuroimaging normative modeling pipelines.

Deep learning methods trained with mean squared error loss are highly sensitive to outliers because the gradient is unbounded. Bounded loss functions (Huber, log-cosh) can help, but they are not standard in current implementations. Pinaya et al. (2019, 2022) reported that variational autoencoder-based normative models are sensitive to training set contamination, with outliers degrading the latent space structure.

### 4.2. Breakdown points and influence functions

The per-method comparison above can be made quantitative through two theoretical concepts from robust statistics (Hampel, 1971; Donoho and Huber, 1983). The breakdown point *ϵ*\* is the smallest fraction of contamination that can drive an estimator to arbitrarily large values; classical estimators have *ϵ*\* = 1*/n* (effectively zero), while the theoretical maximum of 50% is achieved by the median, the Minimum Covariance Determinant (MCD), and Least Trimmed Squares (LTS) (Rousseeuw and Leroy, 1987). The influence function IF(*y*; *T, F*) describes how an infinitesimal contamination at *y* perturbs an estimator *T* at distribution *F*; for OLS it is unbounded (IF *∝ y*), whereas bounded-influence estimators such as the Huber and Tukey bisquare M-estimators have influence functions that saturate or return to zero.

For normative modeling both quantities matter: the breakdown point determines tolerance to contaminated subgroups (e.g., a small site with systematic measurement bias), while the influence function determines sensitivity to individual extreme observations. table 4 and fig. 5 map these properties onto each method.

**Table 4:** Breakdown points and influence function behavior of normative modeling estimators and their robust extensions.

| Method | Breakdown point | Influence function | Reference |
| --- | --- | --- | --- |
| OLS / Residual / ComBat | $1/n \approx 0$ | Unbounded | Hampel (1971) |
| HBR (Gaussian likelihood) | $1/n$ (shrinkage helps) | Unbounded | Kia et al. (2020) |
| HBR (Student-$t$ / SHASH) | Implicitly higher via heavy tails | Bounded | de Boer et al. (2024) |
| GAMLSS (Gaussian) | Low | Unbounded | Stasinopoulos and Rigby (2007) |
| GAMLSS (BCT / BCPE) | Implicitly higher via $t$ tails | Bounded | Rigby and Stasinopoulos (2004) |
| GNM (kernel + log-variance) | Local; depends on bandwidth | Locally bounded | Chen and Bickel (2006); Fan and Gijbels (1996) |
| M-estimator regression | $1/(p+1)$ | Bounded | Maronna et al. (2006) |
| LTS / S-estimator regression | Up to 50% | Bounded | Rousseeuw and Leroy (1987) |
| MM-estimator | 50% with $\sim 95\%$ efficiency | Bounded | Yohai (1987) |
| Kernel + local M-regression | Bandwidth-dependent | Bounded | Härdle (1990) |

**Figure 5:**
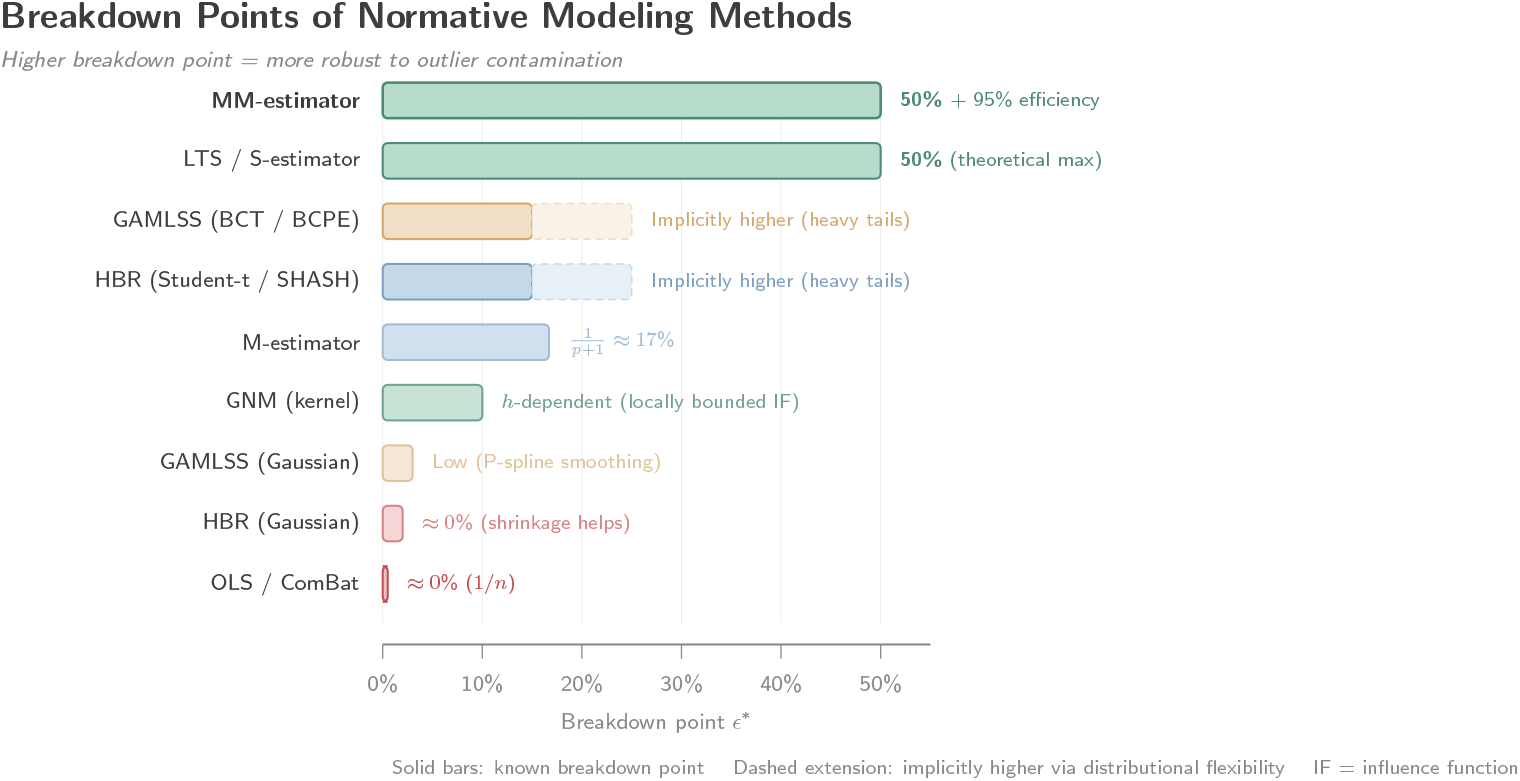
Breakdown points of normative modeling methods and their robust extensions. Classical estimators (OLS, ComBat) have breakdown points near zero; robust alternatives (LTS, MM-estimator) achieve the theoretical maximum of 50%. Dashed extensions indicate methods whose distributional flexibility (heavy-tailed likelihoods) provides implicit resistance beyond the nominal breakdown point.

### 4.3. Robust extensions

None of the off-the-shelf normative modeling methods achieves a high break-down point, which motivates three complementary strategies that can be applied before, during, or after model fitting.

Before fitting, multivariate outlier detection based on the Minimum Covariance Determinant (MCD) estimator (Rousseeuw, 1984; Rousseeuw and Van Driessen, 1999) provides a natural pre-screening step. MCD seeks the *h*-subset of observations whose empirical covariance has minimum determinant, achieving a break-down point of approximately 50% when *h ≈ n/*2. The FAST-MCD algorithm (Rousseeuw and Van Driessen, 1999) and its deterministic variant DetMCD (Hubert et al., 2018) make this computationally feasible for datasets up to ~50,000 subjects. Observations are then screened using robust Mahalanobis distances, with the 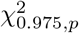 threshold to flag suspected outliers (Rousseeuw and van Zomeren, 1990). For high-dimensional data (*p ≫ n*) where direct MCD is infeasible, a dimensionality reduction step—for instance, *t*-SNE projection (van der Maaten and Hinton, 2008)—precedes the MCD screen; GNM applies this strategy for voxel-wise connectivity data (Bonnelle et al., 2011). Other pre-modeling approaches include Isolation Forest (Liu et al., 1999), which identifies anomalies through recursive random partitioning, Local Outlier Factor (Breunig et al., 2000), which detects density-based local anomalies, and domain-specific tools such as MRIQC (Esteban et al., 2019) that leverage image quality metrics tailored to structural and functional MRI.

During fitting, the most principled approach is to replace the squared loss with a bounded-influence alternative. Classical M-estimators (Huber, 1964; Beaton and Tukey, 1974) use the Huber or Tukey bisquare loss, so that residuals beyond a threshold contribute a constant or zero gradient. MM-estimators (Yohai, 1987) combine the 50% breakdown point of S-estimators in a first stage with the ~95% asymptotic efficiency of a refined M-step in a second stage.

These estimators are readily available in the `robustbase` R package and the `statsmodels` Python library, and could replace OLS in ComBat or the residual model with minimal code changes. Within a Bayesian framework, heavy-tailed likelihoods—Student-*t* for HBR (Geweke, 1993; Lange et al., 1989), SHASH for HBR (de Boer et al., 2024), BCT or BCPE for GAMLSS (Borghi et al., 2006)—achieve the same goal by automatically down-weighting extreme observations through their posterior. These distributional extensions carry negligible computational overhead while substantially reducing sensitivity to heavy-tailed contamination.

After fitting, subjects with extreme *z*-scores (|*z*| *>* 3 or |*z*| *>* 4) can be flagged and removed, but this approach is inherently circular: the *z*-scores themselves are contaminated by the outliers used to fit the model, so the threshold may be miscalibrated (Wolfers et al., 2018). A hybrid strategy—fitting an initial model, removing likely outliers via post-modeling *z*-scores, then refitting on the retained set (Marquand et al., 2014; Rutherford et al., 2022b)—approximates the LTS estimator in spirit and is preferable to a single post-hoc screen. Li et al. (2022) adopted this strategy in the HarMNqEEG project, detecting outliers on the global *z*-scores rather than on the raw data; MCD-based automatic screening can be combined with this step to remove outliers before refitting.

In practice, these three strategies are complementary rather than competing: pre-modeling quality control removes technical artifacts, in-modeling resistance handles residual contamination, and post-modeling calibration checks verify that the resulting *z*-scores are well-behaved. Studies should report their outlier handling explicitly, as most multi-site studies currently underreport removal procedures (Bhagwat et al., 2021), undermining reproducibility.

### 4.4. Summary

table 5 summarizes the native resistance, recommended extensions, and implementation maturity for each method.

**Table 5:** Native resistance to outliers, recommended extensions, and implementation maturity for each normative modeling method.

| Method | Native resistance | Recommended extension | Maturity |
| --- | --- | --- | --- |
| OLS / Residual | None | M-estimator (e.g., **robustbase**) | Mature |
| ComBat | EB shrinkage (across features) | Pre-screening + ComBat-GAM (Pomponio et al., 2020) | Mature |
| HBR | Hierarchical prior | Student-$t$ or SHASH likelihood (de Boer et al., 2024) | Emerging |
| GAMLSS | P-spline smoothing | BCT / BCPE distribution family | Mature |
| GNM | Local kernel averaging | MCD pre-screening + log-space variance | Mature |
| Deep learning | None (with MSE loss) | Huber loss + adversarial training | Early |

## 5. Computational Considerations

As multi-site datasets grow to tens of thousands of subjects with hundreds or thousands of features, computational cost becomes a practical constraint on method selection. The Lifespan Brain Chart Consortium aggregated 101,457 MRI scans (Bethlehem et al., 2022); the UK Biobank imaging extension targets 100,000 subjects (Miller et al., 2016); the ENIGMA consortium aggregates over 50,000 subjects across studies (Thompson et al., 2014). Methods that scale poorly with *N* become infeasible at these sizes.

### 5.1. Asymptotic complexity

table 6 summarizes the per-feature cost of each estimation strategy. Let *N* denote the total number of subjects, *p* the number of fixed-effect covariates, *K* the number of spline basis functions, *B* the number of batches, *M* the kernel grid size per dimension, *d* the covariate dimensionality, *S* the number of MCMC samples, *G* the number of GCV bandwidth evaluations, *M*_ind_ the number of inducing points (sparse GP), and *W* the number of neural network parameters.

**Table 6:** Asymptotic per-feature computational cost of normative modeling methods.

| Method | Training cost | Hyperparameter sel. | Inference cost |
| --- | --- | --- | --- |
| OLS / ComBat | $O(Np^2)$ | Closed form | $O(p)$ |
| GAMLSS (P-spline) | $O(NK^2I)$, $I$ = backfit. iter. | REML / AIC | $O(K)$ |
| HBR (BLR + basis) | $O(Np^2BS)$, MCMC | Post. convergence | $O(pS)$ |
| HBR (exact GP) | $O(N^3)$ | Marginal lik. opt. | $O(N^2)$ |
| HBR (Sparse GP) | $O(NM_{\text{ind}}^2)$ | Inducing pt. opt. | $O(M_{\text{ind}}^2)$ |
| GNM (NUFFT-LPR) | $O((N + M^d \log M)G)$ | GCV (closed form) | $O(M^d)$ |
| Deep learning | $O(NEW)$, $E$ = epochs | Manual / Bayes. opt. | $O(W)$ |

The most consequential distinction is between methods whose training cost scales cubically with *N* (exact GP), quadratically (dense kernel regression without NUFFT), or at most linearly (OLS, GAMLSS backfitting, NUFFT-accelerated kernel regression). For a typical neuroimaging dataset (*N* = 5,000, *m* = 68 ROIs, *d* = 1), all methods except exact GP complete within minutes; for EEG data with thousands of features (*m* = 4,230), the per-feature cost is amortized, but total wall-clock time becomes decisive. fig. 6 illustrates these relationships.

**Figure 6:**
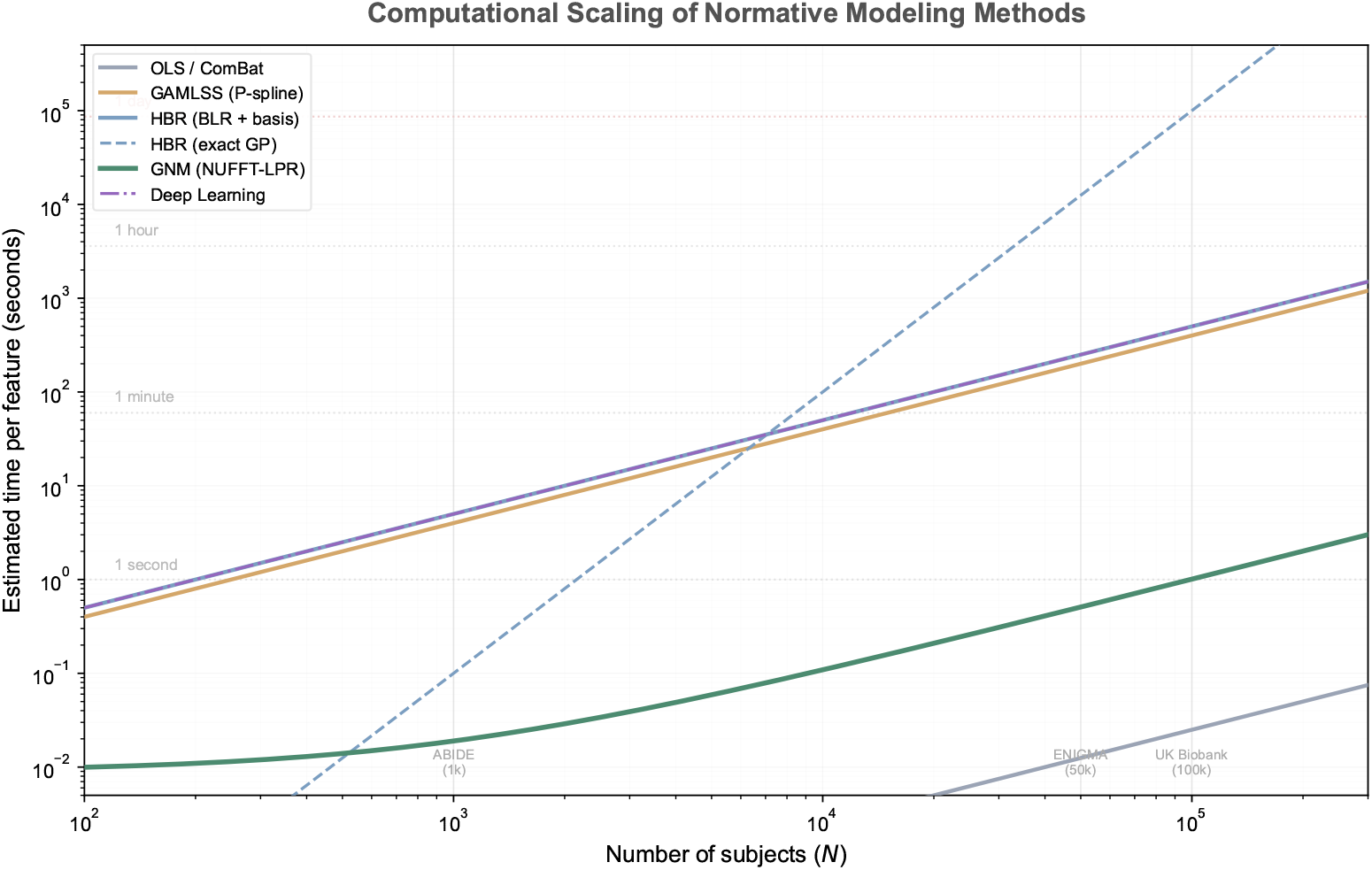
Computational scaling of normative modeling methods. Estimated per-feature training time as a function of sample size *N* on a log–log scale. Reference lines mark practical time limits (1 second, 1 minute, 1 hour, 1 day). Vertical annotations indicate the scale of major neuroimaging consortia.

### 5.2. Empirical scaling and memory

Reported fitting times confirm the theoretical picture, though with important nuances. OLS-based methods (ComBat, ComBat-GAM) are the fastest, completing in seconds to minutes for *N ≤* 10,000 with linear scaling in *N* (Pomponio et al., 2020). GAMLSS with P-splines is nearly as fast for a single covariate—seconds per feature at *N* = 10,000 with convergence in 10–50 backfitting iterations (Stasinopoulos and Rigby, 2007)—but the iterative outer loop for multi-parameter distributions (BCT, BCPE) adds overhead, and the Brain Charts project required days of HPC time for ~120,000 scans across hundreds of features (Bethlehem et al., 2022). HBR with MCMC inference takes minutes to hours per feature at *N ∈* [1,000, 10,000] depending on the basis size and number of chains; variational inference (Kia et al., 2022) reduces this to seconds per feature at the cost of approximation error in the posterior. Exact GP regression becomes impractical beyond *N ≈* 5,000 without sparse approximations (Rasmussen and Williams, 2006), which reduce cost to 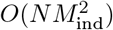 with *M*_ind_ = 100–1,000 inducing points (Quiñonero-Candela and Rasmussen, 2005; Titsias, 2009). NUFFT-accelerated kernel regression runs in milliseconds per feature per bandwidth evaluation at *N* = 5,000 and *M* = 128, with GCV bandwidth selection adding negligible overhead because the Hutchinson trace estimator (Hutchinson, 1990) reuses the same NUFFT pipeline (Li et al., 2026a). Deep learning methods require hours to days of GPU time for training (e.g.,NormVAE: ~6 hours on an A100 for *N* = 35,000 (Lawry Aguila et al., 2022); DeepComBat: comparable GPU time (Hu et al., 2024)), but inference is fast (milliseconds).

Memory requirements follow a parallel hierarchy. OLS and ComBat need only *O*(*Np*) for the design matrix. GAMLSS requires *O*(*NK*) per distribution parameter. HBR with BLR stores the 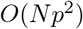 posterior covariance, while exact GP requires the *O*(*N* ^2^) kernel matrix—at *N* = 50,000 this amounts to ~20 GB in double precision, infeasible on standard workstations. Sparse GP and NUFFT-based methods reduce memory to *O*(*NM*_ind_) and *O*(*M* ^*d*^ + *N*) respectively, keeping even large-*N* problems tractable on commodity hardware.

### 5.3. Parallelization and hyperparameter selection

All methods support feature-level parallelization, since each feature is fit independently—an embarrassingly parallel structure that scales linearly with the number of available cores or HPC nodes. MCMC chains in HBR can run in parallel across cores (Gelman et al., 2013), and deep learning naturally leverages GPU acceleration. NUFFT-based methods could benefit from GPU FFT libraries (cuFFT), though this is not yet standard in published pipelines.

The often-overlooked cost of hyperparameter selection can dominate total fitting time. OLS and ComBat have no free hyperparameters beyond the closed-form empirical Bayes shrinkage. GAMLSS integrates smoothing parameter selection into its fitting algorithm via REML or AIC. GP regression requires marginal likelihood optimization, each evaluation costing *O*(*N* ^3^) for exact GP.

Deep learning involves architecture, learning rate, and regularization choices that are typically hand-tuned or selected by Bayesian optimization (Snoek et al., 2012). GCV-based bandwidth selection for kernel regression is efficient because the Hutchinson trace estimator runs within the same NUFFT pass, making bandwidth selection essentially free relative to model fitting.

### 5.4. Practical guidance by dataset size

table 7 translates the above analysis into dataset-size-dependent guidance. For high-dimensional features (*m >* 1000)—typical in voxel-wise analyses or dense connectivity matrices—feature-level parallelization across HPC clusters is necessary regardless of the chosen method.

**Table 7:** Recommended methods by dataset size.

| Dataset size | Recommended methods | Methods to avoid |
| --- | --- | --- |
| Small ($N < 1,000$) | All methods feasible | None |
| Medium ($N \in [1k, 10k]$) | All methods feasible | Exact GP if speed matters |
| Large ($N \in [10k, 100k]$) | ComBat, GAMLSS, GNM, sparse GP, DL | Exact GP |
| Very large ($N > 100k$) | ComBat, GAMLSS, GNM, DL | Exact GP, MCMC HBR |

## 6. Distributed and Federated Estimation

In many clinical settings, raw data cannot leave the originating institution due to privacy regulations (GDPR in Europe, HIPAA in the US, PIPL in China) or data governance agreements. Brain scans are considered sensitive personal data with non-trivial re-identification risks (Schwarz et al., 2019), making federated learning—where computation occurs at each site and only aggregated statistics are shared—a practical necessity. This section analyzes which normative modeling methods admit natural federated decompositions and at what cost.

### 6.1. The federated learning paradigm

The standard federated framework (McMahan et al., 2017; Li et al., 2020) assumes *B* sites each holding a local dataset 𝒟_*b*_ = *{*(**x**_*i*_, *y*_*i*_)*}*_*i∈*site *b*_ of size *N*_*b*_. The protocol alternates between local computation, in which each site derives a summary statistic **s**_*b*_ from 𝒟_*b*_, and central aggregation, in which a server combines *{***s**_1_, …, **s**_*B*_*}* into a global estimate and optionally returns updated parameters for further refinement. A method is *naturally federated* when the local statistics are sufficient for the global estimate—no information is lost by aggregation—and the communication cost is small relative to the data size.

### 6.2. Decomposability across methods

The methods reviewed in section 2 differ substantially in how well they admit such a decomposition, and these differences can be traced directly to the algebraic structure of their estimators.

ComBat is the simplest case. Because the OLS estimates require only the cross-product matrices **X**^*⊤*^**X** and **X**^*⊤*^**y**, which are additive across sites, the global regression can be computed from site-level sufficient statistics without accessing raw data. The batch-specific parameters *γ*_*bv*_ and *δ*_*bv*_ are by definition local to each site. Chen et al. (2022) formalized this as Distributed ComBat and demonstrated on ENIGMA data that it reproduces centralized ComBat with correlations exceeding 0.999, while transmitting only *O*(*p*^2^ + *B*) scalars per feature per round. Bostami et al. (2022) implemented a related decentralized variant within the COINSTAC platform (Plis et al., 2016) using secure multi-party computation. Privacy risk is low because only aggregate statistics cross institutional boundaries.

Hierarchical Bayesian regression also supports a natural—though approximate— federated strategy. Kia et al. (2022) proposed partial pooling, in which each site fits a local Bayesian model and shares its posterior hyperparameters 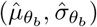 the central server aggregates these into population-level hyperparameters and returns them, analogous to federated variational inference (Corinzia and Buhmann, 2019). The hierarchical prior is well suited to heterogeneous site sizes, as small sites are automatically shrunk more toward the population mean. However, privacy risk is moderate: concentrated posteriors from small samples may leak site-level information, and formal differential privacy guarantees (Dwork et al., 2006) increase posterior uncertainty. Communication cost is *O*(*p*^2^) per feature per round.

GNM has a natural federated structure that follows directly from the additive decomposition of the type-1 nonuniform Fourier transform. The kernel regression sums 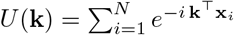 and 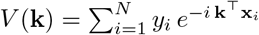 decompose as *U* (**k**) = Σ_*b*_ *U*_*b*_(**k**) and *V* (**k**) = Σ_*b*_ *V*_*b*_(**k**), where each site computes its local Fourier coefficients using the same NUFFT algorithm and transmits only these coefficients—not raw data—to the central server. The server aggregates by summation, computes the global kernel regression on a grid, selects the optimal bandwidth via GCV, and broadcasts the result back to sites for local *z*-score computation. This protocol has several attractive properties. Communication cost is *O*(*M* ^*d*^) complex numbers per feature (e.g., 256 for a one-dimensional covariate with *M* = 128), which is negligible compared to transmitting raw data. Privacy protection is inherent: recovering individual observations from the Fourier coefficients requires solving an underdetermined system when *M* ^*d*^ *≪ N*_*b*_, and calibrated Gaussian noise can provide formal differential privacy guarantees (Dwork et al., 2006). Incremental updates are trivial: when a new site joins, its coefficients are simply added to the existing aggregates, requiring only recomputation of the GCV bandwidth and the grid-domain regression in *O*(*M* ^*d*^ log *M*) time.

GAMLSS currently lacks a federated implementation. The penalized likelihood optimization requires the iterative backfitting algorithm (Rigby and Stasinopoulos, 2005), which updates each distribution parameter sequentially while conditioning on all others—a dependency that resists straightforward decomposition across sites. Developing a federated GAMLSS would require distributed penalized regression, for instance via ADMM (Boyd et al., 2011) or communication-efficient gradient methods, an active research area (Jordan et al., 2019; Fan et al., 2023) that has not yet been applied to normative modeling.

Gaussian process methods pose similar challenges. The kernel matrix **K** couples all observations globally, and the posterior mean at any test point depends on the entire training set. Sparse GP approximations with shared inducing points (Bui et al., 2017; Gal et al., 2014) offer a partial solution: each site computes local sufficient statistics relative to a shared inducing set, and these are aggregated centrally. The inducing point locations, however, may reveal information about each site’s covariate distribution (e.g., its age range), creating a privacy concern. Distributed GP regression has been studied in the machine learning literature (Deisenroth and Ng, 2015; Liu et al., 2020) but has not been applied to normative modeling.

Deep learning methods can leverage federated averaging (FedAvg) (McMahan et al., 2017), in which each site trains locally and shares gradient updates or model weights. However, communication cost scales as *O*(*W*) per round, where *W* is the number of model parameters—often in the millions—and the method is susceptible to catastrophic forgetting when sites have heterogeneous data distributions. Privacy risks are moderate to high, as gradient updates can leak training examples (Zhu et al., 2019).

### 6.3. Out-of-sample batch z-score computation

A central use case of harmonized normative modeling is the assessment of new subjects after the model has been trained—for instance, computing batch-adjusted *z*-scores for patients in a clinical workflow or for subjects acquired from a previously enrolled site without retraining.

The unified equation provides a clean prescription: the batch-specific location and scale estimates 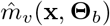 and 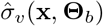obtained during training are reused directly, and the *z*-score is computed 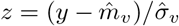. Each method instantiates this prescription differently. In ComBat, the stored batch parameters 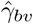 and 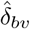are applied to the new observation, followed by a separate normative model for *z*-score computation; ComBat itself does not support out-of-sample harmonization natively, but extensions such as neuroComBat (Fortin et al., 2018) and longitudinal ComBat (Beer et al., 2020) have added this capability. In HBR, the posterior predictive distribution at the new subject’s covariates and batch yields both the mean and variance required for the *z*-score; this is one of the framework’s principal advantages for clinical deployment (Kia et al., 2022). GAMLSS retains the per-site random effect estimates (batch mean and standard deviation) from the training phase, so out-of-sample *z*-scores for subjects from known sites reduce to evaluating the stored fixed and random effect functions at the new covariates (Bethlehem et al., 2022). GNM stores the per-batch mean and standard deviation correction functions 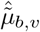and 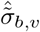estimated during training; a new subject’s batch-adjusted *z*-score is computed directly from these stored batch mean and standard deviation, with no refitting or interpolation required.

Furthermore, by relaxing the definition of “batch” from individual site to a broader grouping—such as scanner manufacturer, acquisition protocol, or geographic region—subjects from nominally new sites can often be assigned to an existing batch category, enabling out-of-sample *z*-score computation using the stored batch parameters without retraining.

### 6.4. Existing platforms

Several platforms support federated or collaborative neuroimaging analyses. COINSTAC (Plis et al., 2016; Gazula et al., 2018) provides a decentralized infrastructure supporting federated regression, ICA, and ComBat through a client-server architecture. Brainlife (Avesani et al., 2019) offers cloud-based reproducible analysis, though data are uploaded to the platform rather than remaining on-site. The Virtual Brain (Sanz-Leon et al., 2013) provides a simulation platform for brain network dynamics that could potentially support federated model fitting. None of these platforms currently support federated kernel regression or NUFFT-based methods; integrating such protocols into existing infrastructure is a natural next step.

### 6.5. Summary

table 8 and fig. 7 summarize the federated readiness of each method. The key distinction is between methods whose estimators decompose into additive site-level statistics (ComBat, GNM) and those that require iterative access to the full dataset (GAMLSS, exact GP). HBR occupies a middle ground, achieving approximate decomposition through hierarchical posterior aggregation.

**Table 8:** Federated readiness of normative modeling methods.

| Method | Federated? | Shared quantity | Privacy risk | Comm. cost | Incremental? |
| --- | --- | --- | --- | --- | --- |
| ComBat | Yes (Chen et al., 2022) | Sufficient statistics | Low | $O(p^2 + B)/\text{feat.}$ | Yes (add batch stats) |
| HBR | Yes (Kia et al., 2022) | Posterior hyperparameters | Medium | $O(p^2)/\text{feat.}$ | Partial (re-aggregate) |
| GAMLSS | No | — | — | — | — |
| GP (sparse) | Difficult | Inducing-point statistics | High | $O(M_{\text{ind}}^2)/\text{feat.}$ | Partial (points may change) |
| GNM (NUFFT) | Yes (natural) | Fourier coefficients | Low | $O(M^d)/\text{feat.}$ | Yes (additive update) |
| Deep learning | Partial (FedAvg) | Gradients / weights | Medium-High | $O(W)/\text{round}$ | Partial (catastrophic forgetting) |

**Figure 7:**
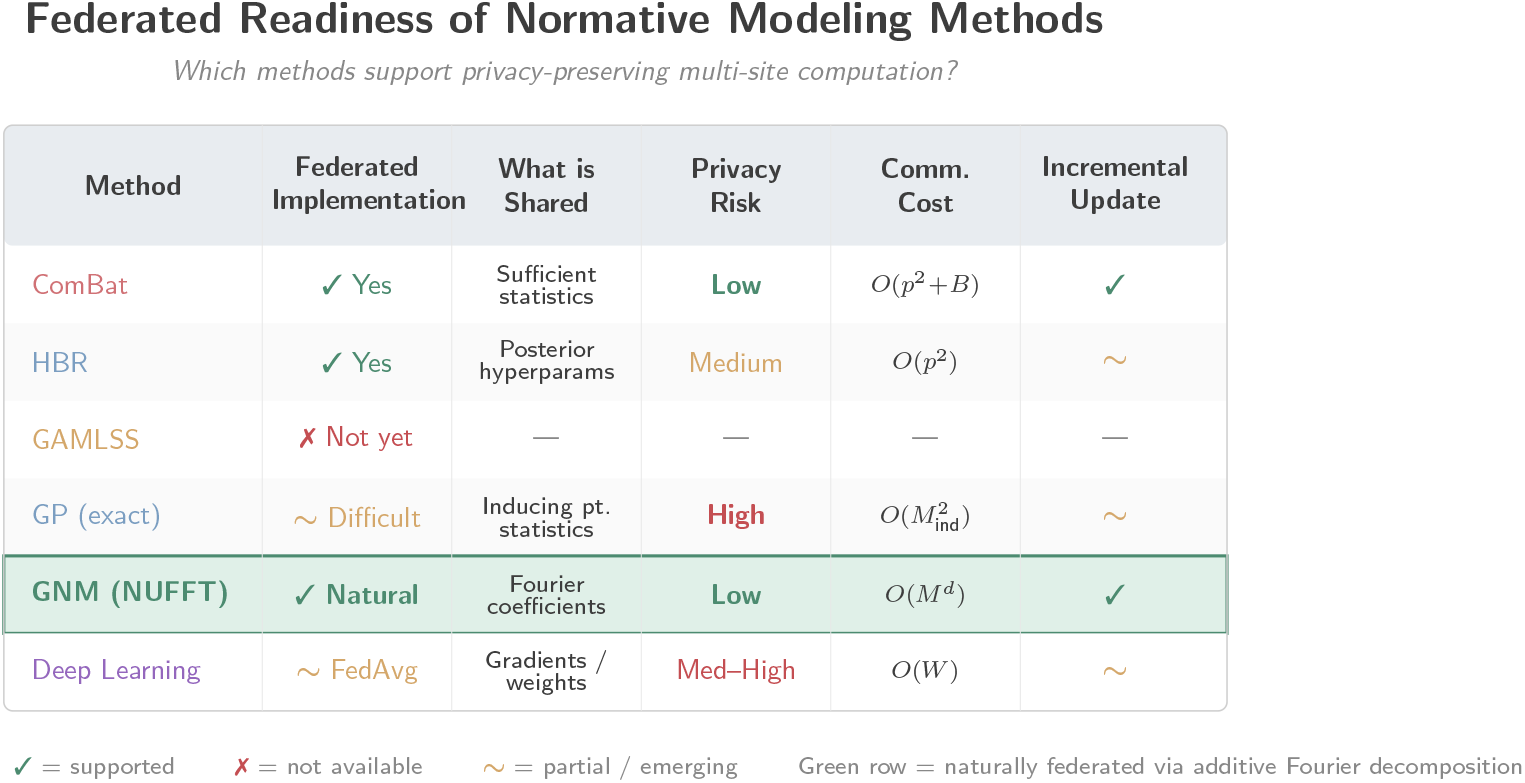
Federated readiness of normative modeling methods. ComBat and GNM support natural federated decomposition with low privacy risk and additive incremental updates. HBR achieves partial pooling through shared posterior hyperparameters. GAMLSS currently lacks a federated implementation due to the sequential dependency of its backfitting algorithm.

## 7. Validation Metrics

Evaluating harmonization and normative modeling methods requires metrics that capture multiple dimensions of performance. A method that removes batch effects perfectly but destroys biological signal is useless; conversely, a method that preserves biology but leaves residual site effects produces misleading *z*-scores. We organize the evaluation framework into three categories: batch effect removal, biological signal preservation, and *z*-score calibration, and argue that for normative modeling the third category should take priority.

### 7.1. Batch effect removal and biological signal preservation

Batch effect removal is typically quantified by the proportion of variance in the harmonized output explained by site. The residual site *R*^2^—computed via ANOVA with site as the sole predictor—should approach zero after harmonization. Fortin et al. (2018) reported reductions from 0.20–0.40 (raw) to 0.01–0.05 (harmonized) for ComBat on cortical thickness, and Pomponio et al. (2020) confirmed similar gains with ComBat-GAM on iSTAGING data. Complementary approaches include training a classifier (random forest, SVM) to predict site from harmonized data, where AUC should equal 0.50 (chance level) (Dinga et al., 2021), and the kernel maximum mean discrepancy (MMD), a nonparametric two-sample statistic that measures distributional distance between sites (Gretton et al., 2012; Dinsdale et al., 2021).

These metrics must, however, be paired with assessments of biological signal preservation. Harmonization that drives site *R*^2^ to zero by also removing age- or sex-related variance is counterproductive. The standard diagnostic is to compare the spatial pattern of age *t*-statistics before and after harmonization: the Spearman correlation *ρ* across features should remain close to one (Fortin et al., 2018; Bethlehem et al., 2022). A more demanding test fits a regression model (ridge regression, SVR) to predict age from harmonized data; when harmonization improves age prediction relative to raw data, the improvement indicates that batch noise was masking biological signal (Li et al., 2026a). Effect size preservation for known group differences (e.g., Cohen’s *d* for sex effects in cortical thickness) provides a third check (Fortin et al., 2018), and for datasets with test–retest scans the intraclass correlation coefficient (ICC) verifies that harmonization does not degrade measurement reliability (Yu et al., 2018; Fortin et al., 2018).

An important subtlety, noted by Li et al. (2026a), is that batch effect metrics computed on harmonized data *y*\* can diverge from those computed on *z*-scores. Methods that jointly estimate the mean and variance (GNM, GAMLSS, HBR) benefit from a self-correcting property: residual site shifts in *y*\* are absorbed during *z*-score construction, so that *R*^2^(site | *z*) can be much lower than *R*^2^(site | *y*\*). Studies that report only data-level metrics may therefore draw misleading conclusions about *z*-score quality.

### 7.2. Z-score calibration

For normative modeling, the *z*-scores are the primary output, and their calibration determines clinical utility. Calibration should be verified along four dimensions.

First, the marginal distribution of *z*-scores in healthy subjects should approximate 𝒩 (0, 1). This is assessed through the overall mean (ideal: 0), standard deviation (ideal: 1), skewness (ideal: 0), and excess kurtosis (ideal: 0), together with the Kolmogorov–Smirnov (KS) test. Fraza et al. (2021) showed that sinharcsinh warping in BLR improved *z*-score normality compared to standard BLR.

Probability–probability and quantile–quantile plots complement these numerical summaries, particularly for assessing tail calibration.

Second, *z*-scores should be independent of covariates. A non-zero Pearson or Spearman correlation between *z* and age, for instance, indicates that the normative model has not fully captured the age trajectory, leaving systematic age-dependent bias. Stratified calibration—computing *z*-score mean and variance within age bins—should yield constant values across bins.

Third, *z*-score distributions should be identical across sites. Per-site means 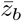 and standard deviations *s*_*b*_ should be constant, verifiable through Levene’s test for variance equality and pairwise KS tests.

Fourth, calibration in the tails deserves particular attention because clinical decisions are made at thresholds such as |*z*| *>* 1.96 or |*z*| *>* 2.58. If the observed proportion of healthy subjects exceeding the threshold departs substantially from the expected 5% or 1%, the *z*-scores are miscalibrated precisely where they matter most. de Boer et al. (2024) showed that SHASH-HBR improved tail calibration compared to Gaussian HBR, reducing the excess of extreme *z*-scores in brain regions with skewed distributions.

When *z*-scores are used for clinical applications, their utility can be assessed through group discrimination (Cohen’s *d* and AUC between patients and controls), individual-level prediction of clinical outcomes (Marquand et al., 2014), and extreme deviation analysis—counting the number of brain regions with |*z*| *>* 2 per subject and comparing the distribution in healthy subjects against the expected binomial with *p* = 0.05 (Wolfers et al., 2018; Zabihi et al., 2019).

### 7.3. Published benchmark comparisons

table 9 collects published performance metrics across methods. Because datasets, preprocessing, and evaluation protocols differ across studies, direct comparison across rows should be made cautiously. Nonetheless, the table reveals a general pattern: methods that jointly model the location and scale (GNM, GAMLSS, HBR) tend to achieve lower residual *R*^2^ at the *z*-score level than two-step approaches (ComBat followed by a separate normative model),consistent with the self-correcting property discussed above.

**Table 9:** Published performance metrics for harmonization and normative modeling methods. CT = cortical thickness. Values are reported as given in the original sources; cross-row comparisons should account for dataset and protocol differences.

| Method | Dataset | $N$ | Metric | Value | Source |
| --- | --- | --- | --- | --- | --- |
| ComBat | ABIDE (CT) | 1,112 | $R^2(\text{site} \mid y^*)$ | 0.01–0.05 | Fortin et al. (2018) |
| ComBat | iSTAGING (CT) | 10,202 | Age $\rho$ preservation | 0.97 | Pomponio et al. (2020) |
| ComBat-GAM | iSTAGING (CT) | 10,202 | Age prediction $R^2$ | 0.82 | Pomponio et al. (2020) |
| GAMLSS | Brain Charts (CT) | 123,984 | KS test ($z$ normality) | Improved vs. Gaussian | Bethlehem et al. (2022) |
| HBR | Multi-site (CT) | 7,286 | $R^2(\text{site} \mid z)$ | $< 0.01$ | Kia et al. (2022) |
| HBR-SHASH | OpenNeuro (CT) | $\sim 5,000$ | Tail calibration | Improved vs. Gaussian HBR | de Boer et al. (2024) |
| Warped BLR | Multi-site (CT) | 7,286 | KS test ($z$ normality) | Improved vs. BLR | Fraza et al. (2021) |
| GNM | ABIDE (CT) | 387 | $R^2(\text{site} \mid z)$ | 0.0000 | Li et al. (2026a) |
| GNM | ABIDE (CT) | 387 | Cross-site mean SD | 0.0000 | Li et al. (2026a) |
| GNM | ABIDE (CT) | 387 | Age prediction RMSE (yr) | 1.86 | Li et al. (2026a) |
| GNM | ABIDE (CT) | 387 | Age $t$-stat $\rho$ | 0.88 | Li et al. (2026a) |
| GNM | HarMNqEEG (EEG) | 1,564 | Cross-site mean SD | 0.0007 | Li et al. (2026a) |
| GNM | HarMNqEEG (EEG) | 1,564 | $R^2(\text{site} \mid z)$ | 0.000014 | Li et al. (2026a) |
| GNM | HarMNqEEG (EEG) | 1,564 | Sig. site channels (Bonf.) | 0/18 | Li et al. (2026a) |
| ComBat+Norm | ABIDE (CT) | 387 | $R^2(\text{site} \mid z)$ | 0.0061 | Li et al. (2026a) |
| GAMLSS | ABIDE (CT) | 387 | $R^2(\text{site} \mid z)$ | 0.0004 | Li et al. (2026a) |
| DeepComBat | Multi-site (MRI) | $\sim 3,000$ | Site prediction AUC | $\sim 0.55$ | Hu et al. (2024) |
| Adversarial | Multi-site (MRI) | $\sim 5,000$ | MMD | Reduced vs. raw | Dinsdale et al. (2021) |

### 7.4. Recommendations for evaluation

Based on the literature reviewed above, we offer several recommendations. Studies should report metrics from all three categories—batch removal, biological preservation, and *z*-score calibration—because reporting only one category provides an incomplete and potentially misleading picture. When the goal is normative modeling, *z*-score-level metrics should take priority over data-level metrics. Tail calibration deserves explicit attention whenever *z*-scores will be used for clinical thresholding. Evaluations should span multiple datasets covering different modalities, sample sizes, and numbers of sites to assess generalizability. Computational cost—wall-clock time and memory—should be reported to enable practical method selection. Code and data availability remain indispensable for reproducibility; the PCNtoolkit (Rutherford et al., 2022a), the `gamlss` R package (Stasinopoulos and Rigby, 2007; Stasinopoulos et al., 2017), and the GNM-ToolBox (Li et al., 2026a) are examples of open-source implementations that facilitate direct comparison.

## 8. Open Challenges and Future Directions

### 8.1. Scope and limitations of the unified equation

The unified location–scale equation (eq. (6)) provides a common language for comparing LMM-based harmonization methods, but its scope is deliberately limited. Methods that do not expose an explicit location–scale pair—deep learning approaches such as DeepComBat, adversarial unlearning, and conditional VAEs (section 3.5)—fall outside the framework. As neural network–based methods mature, the proportion of the field’s methodological toolkit that sits outside the unified equation will grow. The framework is therefore best understood as a lens for a particular (albeit currently dominant) family of methods, not as an exhaustive taxonomy.

A second limitation concerns the Gaussian assumption on *ε*. While the unified equation accommodates heteroscedasticity through the scale function *σ*_*v*_(**x, Θ**), the *z*-score interpretation assumes that the standardized residuals are at least approximately Gaussian. Extensions to non-Gaussian likelihoods— Student-*t*, SHASH, BCT—relax this assumption within each method, but the unified equation itself does not prescribe how to handle skewness or heavy tails, leaving this choice to the practitioner.

### 8.2. When harmonization may be harmful

The framework implicitly assumes that batch effects are purely nuisance variation that should be removed. This assumption can fail when batch effects interact with biology—for instance, when a scanner’s sensitivity to cortical thinning varies with age, or when site-level demographic differences (socioeconomic status, ethnicity) are confounded with scanner identity. In such cases, additive or multiplicative harmonization may remove genuine biological signal along with the batch effect. Causal approaches to harmonization, using directed acyclic graphs (DAGs) to distinguish confounding from mediation, could provide principled criteria for what should and should not be removed. This direction is particularly relevant when the goal extends beyond *z*-score computation to causal inference about brain–behavior relationships. At a minimum, practitioners should verify that known biological effects (age, sex) are preserved after harmonization and consider whether site-biology confounding is plausible in their study design.

### 8.3. Multivariate norms and spatial covariance

Current methods fit each feature independently, ignoring the spatial covariance structure across brain regions. A multivariate normative model that accounts for cross-region correlations could improve both normative trajectory estimates and batch effect estimates by borrowing strength across features. The empirical Bayes shrinkage employed by ComBat across features represents a simple form of such borrowing, but a full multivariate extension—combining nonparametric regression with spatial covariance modeling—remains an open problem. Methods such as NormVAE (Lawry Aguila et al., 2022) and DeepComBat (Hu et al., 2024) operate in a multivariate latent space but sacrifice the explicit distributional framework needed for calibrated *z*-scores.

### 8.4. Non-Euclidean descriptive parameters

Many brain-derived measures do not reside in Euclidean space. Connectivity matrices—correlations, covariances—are symmetric positive definite (SPD) matrices that lie on a Riemannian manifold. Applying standard *z*-score transformations to vectorized connectivity matrices ignores the geometric structure and can produce nonsensical results. Riemannian mapping techniques, such as log-Euclidean or affine-invariant metrics, can transform SPD matrices into a tangent space where Euclidean methods apply, but integrating these transformations into the unified normative modeling framework is largely unexplored.

### 8.5. Continual and online normative modeling

As new sites join a consortium or existing sites acquire new data, the normative model must be updated. Most current methods require refitting from scratch, which is computationally expensive and requires access to all historical data. Online or continual learning approaches, where the model is updated incrementally with new data, would be valuable for large-scale, evolving consortia. The federated NUFFT structure of GNM (section 6) is naturally suited to incremental updates, as new sites contribute their Fourier coefficients to the existing aggregates, but analogous incremental strategies for GAMLSS and HBR remain undeveloped.

### 8.6. Verification of batch effects

A basic but often overlooked question is: what constitutes a “batch effect”? Current methods assume that site is the primary source of non-biological variability, but in practice batch effects may arise from scanner manufacturer, acquisition protocol, software version, operator, time of day, or combinations thereof. No existing harmonization method supports systematic verification of which factors contribute to batch effects. Developing diagnostic tools that decompose the total non-biological variance into attributable components would help researchers make informed decisions about what to harmonize.

## 9. Conclusion

The rapid expansion of multi-center neuroimaging consortia has made batch-harmonized normative modeling indispensable, yet the field has lacked a common theoretical framework for understanding how existing harmonization methods support the computation of site-invariant *z*-scores. This review addresses that gap.

We have shown that all widely used LMM-based harmonization methods— the residual model, ComBat, GAMLSS, HBR, and GNM—are special cases of a single location–scale equation:

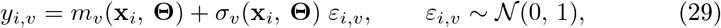

which we term the unified form of batch harmonization equation for normative modeling. The methods differ along three dimensions: the functional forms of the location *m*_*v*_ and scale *σ*_*v*_ (linear, additive spline, Bayesian nonparametric,or kernel); how batch effects enter **Θ** (constant, linear, hierarchical prior, or functional); and the estimation strategy (OLS, penalized likelihood, MCMC, or GCV with NUFFT). This unified perspective reveals that the choice among methods is not about different modeling philosophies, but about the flexibility–computation trade-off along a spectrum from fully parametric (ComBat) to fully nonparametric (GNM).

Building on this framework, we have compared the regression engines underlying each method, quantified their sensitivity to outliers through breakdown points and influence functions, evaluated computational scaling from hundreds to hundreds of thousands of subjects, analyzed federated decomposability for privacy-preserving multi-center computation, and clarified how each method supports out-of-sample batch-adjusted *z*-score computation using stored batch parameters.

By providing a common theoretical language that makes assumptions explicit and trade-offs transparent, the unified equation establishes a principled foundation for method selection and paves the way toward reliable, scalable, and privacy-aware normative modeling across the rapidly expanding landscape of multi-center neuroimaging.

## CRediT authorship contribution statement

**Min Li**: Conceptualization, Investigation, Formal analysis, Visualization, Writing – original draft. **Ying Wang**: Investigation, Visualization, Writing – review & editing. **Yajun Shen**: Investigation, Writing – review & editing. **Lin An**: Writing – review & editing. **Gangyong Jia**: Writing – review & editing. **Maria L. Bringas-Vega**: Writing – review & editing. **Pedro Antonio Valdés-Sosa**: Writing – review & editing.

## Data and code availability

The Generalized Normative Modeling toolbox is available at https://github.com/LMNonlinear/Generalized-Normative-Modeling. The fastLPR library is available at https://github.com/rigelfalcon/fastLPR. The PCN-toolkit for HBR is available at https://github.com/amarquand/PCNtoolkit. The gamlss R package is available from CRAN. No new data were generated for this review; all reported results are from the cited publications.

## Acknowledgments

M.L. was supported by Hangzhou Dianzi University Seed Fund Project (KYS055623037) and Zhejiang Provincial Higher Education Institutions’ Basic Operations Project (GK239909299001-025). G.J. was supported by the Zhejiang Key Research and Development Program under Grant No. 2026C01028 and No. 2023C03194. P.A.V.-S. was supported by National Key R&D Program of China (2024YFE0215100), the CNS Program of the University of Electronic Science and Technology of China (UESTC) (Grant No. Y03023206100204).

## References

Avesani, P., McPherson, B., Hayashi, S., Caiafa, C.F., Henschel, R., Garyfallidis, E., Kitchell, L., Bullock, D., Patterson, A., Olivetti, E., Sporns, O., Saykin, A.J., Wang, L., Dinov, I., Pestilli, F., 2019. The open diffusion data derivatives, brain data upcycling via integrated publishing of derivatives and reproducible open cloud services. Scientific Data 6, 69. doi:10.1038/s41597-019-0073-y.

Bayer, J.M.M., Dinga, R., Kia, S.M., Kottaram, A.R., Wolfers, T., Lv, J., Zalesky, A., Schmaal, L., Marquand, A.F., 2022. Accommodating site variation in neuroimaging data using normative and hierarchical Bayesian models. NeuroImage 264, 119699. doi:10.1016/j.neuroimage.2022.119699.

Beaton, A.E., Tukey, J.W., 1974. The fitting of power series, meaning polynomials, illustrated on band-spectroscopic data. Technometrics 16, 147–185. doi:10.1080/00401706.1974.10489171.

Beer, J.C., Tustison, N.J., Cook, P.A., Davatzikos, C., Sheline, Y.I., Shinohara, R.T., Linn, K.A., 2020. Longitudinal ComBat: A method for harmonizing longitudinal multi-scanner imaging data. NeuroImage 220, 117129. doi:10.1016/j.neuroimage.2020.117129.

Bethlehem, R.A.I., Seidlitz, J., White, S.R., Vogel, J.W., Anderson, K.M., Adamson, C., Adler, S., Alfaro-Almagro, F., Anderson-Davies, S., Avelar-Pereira, B., et al., 2022. Brain charts for the human lifespan. Nature 604, 525–533. doi:10.1038/s41586-022-04554-y.

Bhagwat, N., Barry, A., Bhatt, P., Bhattasali, S., Bhogawar, M., Bond, M., et al., 2021. Understanding the impact of preprocessing pipelines on neuroimaging cortical surface analyses. GigaScience 10, giaa155. doi:10.1093/gigascience/giaa155.

de Boer, A.A.A., Kia, S.M., Rutherford, S., Zabihi, M., Fraza, C., Barkema, P., Westlye, L.T., Andreassen, O.A., Hinne, M., Beckmann, C.F., Marquand, A.F., 2024. Non-Gaussian normative modelling with hierarchical Bayesian regression. Imaging Neuroscience 2, 1–22. doi:10.1162/imag_a_00132.

Bonnelle, V., Leech, R., Kinnunen, K.M., Ham, T.E., Beckmann, C.F., De Boissezon, X., Greenwood, R.J., Sharp, D.J., 2011. Default mode network connectivity predicts sustained attention deficits after traumatic brain injury. Journal of Neuroscience 31, 13442–13451. doi:10.1523/JNEUROSCI.1163-11.2011.

Borghi, E., de Onis, M., Garza, C., Van den Broeck, J., Frongillo, E.A., Grummer-Strawn, L., Van Buuren, S., Pan, H., Molinari, L., Martorell, R., et al., 2006. Construction of the World Health Organization child growth standards: selection of methods for attained growth curves. Statistics in Medicine 25, 247–265. doi:10.1002/sim.2227.

Bostami, B., Hillary, F.G., van der Horn, H.J., van der Naalt, J., Calhoun, V.D., Vergara, V.M., 2022. A decentralized ComBat algorithm and applications to functional network connectivity. Frontiers in Neurology 13, 826734. doi:10.3389/fneur.2022.826734.

Boyd, S., Parikh, N., Chu, E., Peleato, B., Eckstein, J., 2011. Distributed optimization and statistical learning via the alternating direction method of multipliers. Foundations and Trends in Machine Learning 3, 1–122. doi:10.1561/2200000016.

Breunig, M.M., Kriegel, H.P., Ng, R.T., Sander, J., 2000. LOF: Identifying density-based local outliers, in: Proceedings of the 2000 ACM SIGMOD International Conference on Management of Data, pp. 93–104. doi:10.1145/335191.335388.

Bui, T.D., Yan, J., Turner, R.E., 2017. A unifying framework for Gaussian process pseudo-point approximations using power expectation propagation. Journal of Machine Learning Research 18, 1–72.

Chen, A., Bickel, P.J., 2006. Efficient independent component analysis. Annals of Statistics 34, 2825–2855. doi:10.1214/009053606000000939.

Chen, A.A., Luo, C., Chen, Y., Shinohara, R.T., Shou, H., 2022. Privacy-preserving harmonization via distributed ComBat. NeuroImage 248, 118822. doi:10.1016/j.neuroimage.2021.118822.

Cole, J.H., Franke, K., 2017. Predicting age using neuroimaging: innovative brain ageing biomarkers. Trends in Neurosciences 40, 681–690. doi:10.1016/j.tins.2017.10.001.

Cole, J.H., Poudel, R.P.K., Tsagkrasoulis, D., Caan, M.W.A., Steves, C., Spector, T.D., Montana, G., 2017. Predicting brain age with deep learning from raw imaging data results in a reliable and heritable biomarker. NeuroImage 163, 115–124. doi:10.1016/j.neuroimage.2017.07.059.

Cole, T.J., 2012. The development of growth references and growth charts. Annals of Human Biology 39, 382–394. doi:10.3109/03014460.2012.694475.

Corinzia, L., Buhmann, J.M., 2019. Variational federated multi-task learning, in: NeurIPS Workshop on Federated Learning.

Deisenroth, M.P., Ng, J.W., 2015. Distributed Gaussian processes, in: Proceedings of the 32nd International Conference on Machine Learning, pp. 1481–1490.

Di Martino, A., Yan, C.G., Li, Q., Denio, E., Castellanos, F.X., Alaerts, K., Anderson, J.S., Assaf, M., Bookheimer, S.Y., Dapretto, M., et al., 2014. The autism brain imaging data exchange: towards a large-scale evaluation of the intrinsic brain architecture in autism. Molecular Psychiatry 19, 659–667. doi:10.1038/mp.2013.78.

Dinga, R., Fraza, C.J., Bayer, J.M.M., Kia, S.M., Penninx, B.W.J.H., Veltman, D.J., Marquand, A.F., 2021. Normative modeling of neuroimaging data using generalized additive models of location scale and shape. bioRxiv doi:10.1101/2021.06.14.448106.

Dinsdale, N.K., Jenkinson, M., Namburete, A.I.L., 2021. Deep learning-based unlearning of dataset bias for MRI harmonisation and confound removal. NeuroImage 228, 117689. doi:10.1016/j.neuroimage.2020.117689.

Donoho, D.L., Huber, P.J., 1983. The notion of breakdown point, in: A Festschrift for Erich L. Lehmann. Wadsworth, pp. 157–184.

Dwork, C., McSherry, F., Nissim, K., Smith, A., 2006. Calibrating noise to sensitivity in private data analysis, in: Theory of Cryptography Conference, pp. 265–284. doi:10.1007/11681878_14.

Eilers, P.H.C., Marx, B.D., 1996. Flexible smoothing with B-splines and penalties. Statistical Science 11, 89–121. doi:10.1214/ss/1038425655.

Esteban, O., Markiewicz, C.J., Blair, R.W., Moodie, C.A., Isik, A.I., Erramuzpe, A., Kent, J.D., Goncalves, M., DuPre, E., Snyder, M., Oya, H., Ghosh, S.S., Wright, J., Durnez, J., Poldrack, R.A., Gorgolewski, K.J., 2019. fMRIPrep: a robust preprocessing pipeline for functional MRI. Nature Methods 16, 111–116. doi:10.1038/s41592-018-0235-4.

Fan, J., 1993. Local linear regression smoothers and their minimax efficiencies. Annals of Statistics 21, 196–216. doi:10.1214/aos/1176349022.

Fan, J., Gijbels, I., 1996. Local Polynomial Modelling and Its Applications. Chapman and Hall, London. doi:10.1201/9780203748725.

Fan, J., Guo, Y., Wang, K., 2023. Communication-efficient accurate statistical estimation. Journal of the American Statistical Association 118, 1000–1010. doi:10.1080/01621459.2021.1969238.

Fernandez, C., Steel, M.F.J., 1998. On Bayesian modeling of fat tails and skewness. Journal of the American Statistical Association 93, 359–371. doi:10.1080/01621459.1998.10474117.

Fortin, J.P., Cullen, N., Sheline, Y.I., Taylor, W.D., Aselcioglu, I., Cook, P.A.,Adams, P., Cooper, C., Fava, M., McGrath, P.J., McInnis, M., Phillips, M.L., Trivedi, M.H., Weissman, M.M., Shinohara, R.T., 2018. Harmonization of cortical thickness measurements across scanners and sites. NeuroImage 167, 104–120. doi:10.1016/j.neuroimage.2017.11.024.

Fraza, C.J., Dinga, R., Beckmann, C.F., Marquand, A.F., 2021. Warped Bayesian linear regression for normative modelling of big data. NeuroImage 245, 118715. doi:10.1016/j.neuroimage.2021.118715.

Gal, Y., van der Wilk, M., Rasmussen, C.E., 2014. Distributed variational inference in sparse Gaussian process regression and latent variable models, in: Advances in Neural Information Processing Systems.

Gazula, H., Baker, B.T., Damaraju, E., Plis, S.M., Panta, S.R., Silva, R.F., Calhoun, V.D., 2018. Decentralized analysis of brain imaging data: voxel-based morphometry and dynamic functional network connectivity. Frontiers in Neuroinformatics 12, 55. doi:10.3389/fninf.2018.00055.

Gelman, A., Carlin, J.B., Stern, H.S., Dunson, D.B., Vehtari, A., Rubin, D.B., 2013. Bayesian Data Analysis. 3rd ed., Chapman and Hall/CRC. doi:10.1201/b16018.

Geweke, J., 1993. Bayesian treatment of the independent Student-*t* linear model. Journal of Applied Econometrics 8, S19–S40. doi:10.1002/jae.3950080504.

Greengard, L., Lee, J.Y., 2004. Accelerating the nonuniform fast Fourier transform. SIAM Review 46, 443–454. doi:10.1137/S003614450343200X.

Gretton, A., Borgwardt, K.M., Rasch, M.J., Schölkopf, B., Smola, A., 2012. A kernel two-sample test. Journal of Machine Learning Research 13, 723–773.

Hampel, F.R., 1971. A general qualitative definition of robustness. Annals of Mathematical Statistics 42, 1887–1896. doi:10.1214/aoms/1177693054.

Härdle, W., 1990. Applied Nonparametric Regression. Cambridge University Press. doi:10.1017/CCOL0521382483.

Horng, H., Singh, A., Yousefi, B., Cohen, E.A., Haghighi, B., Shinohara, R.T., Madabhushi, A., 2022. Generalized ComBat harmonization methods for radiomic features with multi-modal distributions and multiple batch effects. Scientific Reports 12, 4493. doi:10.1038/s41598-022-08412-9.

Hu, F., Lucas, A., Chen, A.A., Coleman, K., Horng, H., Ng, R.W.S., Tustison, N.J., Davis, K.A., Shou, H., Li, M., Shinohara, R.T., 2024. DeepComBat: A statistically motivated, hyperparameter-robust, deep learning approach to harmonization of neuroimaging data. Human Brain Mapping 45, e26708. doi:10.1002/hbm.26708.

Huber, P.J., 1964. Robust estimation of a location parameter. Annals of Mathematical Statistics 35, 73–101. doi:10.1214/aoms/1177703732.

Hubert, M., Debruyne, M., Rousseeuw, P.J., 2018. Minimum covariance determinant and extensions. Wiley Interdisciplinary Reviews: Computational Statistics 10, e1421. doi:10.1002/wics.1421.

Hutchinson, M.F., 1990. A stochastic estimator of the trace of the influence matrix for Laplacian smoothing splines. Communications in Statistics – Simulation and Computation 19, 433–450. doi:10.1080/03610919008812866.

Johnson, W.E., Li, C., Rabinovic, A., 2007. Adjusting batch effects in microarray expression data using empirical Bayes methods. Biostatistics 8, 118–127. doi:10.1093/biostatistics/kxj037.

Jordan, M.I., Lee, J.D., Yang, Y., 2019. Communication-efficient distributed statistical inference. Journal of the American Statistical Association 114, 668–681. doi:10.1080/01621459.2018.1429274.

Kia, S.M., Huijsdens, H., Dinga, R., Wolfers, T., Mennes, M., Andreassen, O.A., Westlye, L.T., Beckmann, C.F., Marquand, A.F., 2020. Hierarchical Bayesian regression for multi-site normative modeling of neuroimaging data, in: Medical Image Computing and Computer Assisted Intervention – MICCAI 2020, Springer. pp. 699–709. doi:10.1007/978-3-030-59728-3_68.

Kia, S.M., Huijsdens, H., Rutherford, S., de Boer, A., Dinga, R., Wolfers, T., Berthet, P., Mennes, M., Andreassen, O.A., Westlye, L.T., Beckmann, C.F., Marquand, A.F., 2022. Closing the life-cycle of normative modeling using federated hierarchical Bayesian regression. PLoS ONE 17, e0278776. doi:10.1371/journal.pone.0278776.

Lange, K.L., Little, R.J.A., Taylor, J.M.G., 1989. Robust statistical modeling using the *t* distribution. Journal of the American Statistical Association 84, 881–896. doi:10.1080/01621459.1989.10478852.

Lawry Aguila, A., Chapman, J., Janahi, M., Altmann, A., 2022. Conditional VAEs for confound removal and normative modelling of neurodegenerative diseases, 430–440 doi:10.1007/978-3-031-16431-6_41.

Li, M., Wang, Y., Chen, Y., Dubois, A.E.E., Jia, G., Wu, Q., Bringas-Vega, M.L., Dumas, G., Valdés-Sosa, P.A., 2025. Aperiodic and periodic EEG component lifespan trajectories: Monotonic decrease versus growth-then-decline. bioRxiv doi:10.1101/2025.08.26.672407.

Li, M., Wang, Y., Lopez-Naranjo, C., Abd Hamid, A.I., Evans, A.C., Savostyanov, A.N., Calzada-Reyes, A., Areces-Gonzalez, A., Villringer, A., Tobon-Quintero, C.A., Garcia-Agustin, D., Paz-Linares, D., Yao, D., Dong, L., Aubert-Vazquez, E., Reza, F., Omar, H., Abdullah, J.M., Galler, J.R., Ochoa-Gomez, J.F., Prichep, L.S., Galan-Garcia, L., Morales-Chacon, L., Valdes-Sosa, M.J., Tröndle, M., Mohd Zulkifly, M.F.B., Abdul Rahman, M.R.B., Milakhina, N.S., Langer, N., Rudych, P., Hu, S., Koenig, T., Virues-Alba, T.A., Lei, X., Bringas-Vega, M.L., Bosch-Bayard, J.F., Valdes-Sosa, P.A., 2022. Harmonized-multinational qEEG norms (HarMNqEEG). NeuroImage 256, 119190. doi:10.1016/j.neuroimage.2022.119190.

Li, M., Wang, Y., Shen, Y., Bringas-Vega, M.L., Valdés-Sosa, P.A., An, L., Jia, G., 2026a. Generalized normative modeling: A one-step hierarchical kernel framework for multi-site brain charts with self-correcting z-scores. NeuroImage In preparation.

Li, M., Wang, Y., Shen, Y., Bringas-Vega, M.L., Valdés-Sosa, P.A., An, L., Jia, G., 2026b. Regression algorithms in normative modeling for neuroimaging: A methodological review In preparation.

Li, T., Sahu, A.K., Zaheer, M., Sanjabi, M., Talwalkar, A., Smith, V., 2020. Federated optimization in heterogeneous networks, in: Proceedings of Machine Learning and Systems (MLSys), pp. 429–450.

Liu, H., Ong, Y.S., Shen, X., Cai, J., 2020. When Gaussian process meets big data: A review of scalable GPs. IEEE Transactions on Neural Networks and Learning Systems 31, 4405–4423. doi:10.1109/TNNLS.2019.2957109.

Liu, R.Y., Parelius, J.M., Singh, K., 1999. Multivariate analysis by data depth:descriptive statistics, graphics and inference. Annals of Statistics 27, 783–858. doi:10.1214/aos/1018031260.

van der Maaten, L., Hinton, G., 2008. Visualizing data using t-SNE. Journal of Machine Learning Research 9, 2579–2605.

Maronna, R.A., Martin, R.D., Yohai, V.J., 2006. Robust Statistics: Theory and Methods. John Wiley & Sons. doi:10.1002/0470010940.

Marquand, A.F., Brammer, M., Williams, S.C.R., Doyle, O.M., 2014. Bayesian multi-task learning for decoding multi-subject neuroimaging data. NeuroImage 92, 298–311. doi:10.1016/j.neuroimage.2014.02.008.

Marquand, A.F., Kia, S.M., Zabihi, M., Wolfers, T., Buitelaar, J.K., Beckmann, C.F., 2019. Conceptualizing mental disorders as deviations from normative functioning. Molecular Psychiatry 24, 1415–1424. doi:10.1038/s41380-019-0441-1.

Marquand, A.F., Rezek, I., Buitelaar, J.K., Beckmann, C.F., 2016. Understanding heterogeneity in clinical cohorts using normative models: beyond case-control studies. Biological Psychiatry 80, 552–561. doi:10.1016/j.biopsych.2015.12.023.

McMahan, B., Moore, E., Ramage, D., Hampson, S., Arcas, B.A.y., 2017. Communication-efficient learning of deep networks from decentralized data, in: Proceedings of the 20th International Conference on Artificial Intelligence and Statistics, pp. 1273–1282.

Miller, K.L., Alfaro-Almagro, F., Bangerter, N.K., Thomas, D.L., Yacoub, E., Xu, J., Bartsch, A.J., Jbabdi, S., Sotiropoulos, S.N., Andersson, J.L.R., Griffanti, L., Douaud, G., Okell, T.W., Weale, P., Dragonu, I., Garas, S., Hudson, N., Collins, R., Jenkinson, M., Matthews, P.M., Smith, S.M., 2016. Multi-modal population brain imaging in the UK Biobank prospective epidemiological study. Nature Neuroscience 19, 1523–1536. doi:10.1038/nn.4393.

Moyer, D., Ver Steeg, G., Tax, C.M.W., Thompson, P.M., 2020. Scanner invariant representations for diffusion MRI harmonization. Magnetic Resonance in Medicine 84, 2174–2189. doi:10.1002/mrm.28243.

Pinaya, W.H.L., Mechelli, A., Sato, J.R., 2019. Using deep autoencoders to identify abnormal brain structural patterns in neuropsychiatric disorders: a large-scale multi-sample study. Human Brain Mapping 40, 944–954. doi:10.1002/hbm.24423.

Pinaya, W.H.L., Tudosiu, P.D., Gray, R., Rees, G., Nachev, P., Ourselin, S., Cardoso, M.J., 2022. Unsupervised brain imaging anomaly detection and segmentation with transformers. Medical Image Analysis 79, 102475. doi:10.1016/j.media.2022.102475.

Plis, S.M., Sarwate, A.D., Wood, D., Dieringer, C., Landis, D., Reed, C., Panta, S.R., Turner, J.A., Shoemaker, J.M., Carter, K.W., Thompson, P., Hutchison, K., Calhoun, V.D., 2016. COINSTAC: A privacy enabled model and prototype for leveraging and processing decentralized brain imaging data. Frontiers in Neuroscience 10, 365. doi:10.3389/fnins.2016.00365.

Pomponio, R., Erus, G., Habes, M., Doshi, J., Srinivasan, D., Mamourian, E., Bashyam, V., Nasrallah, I.M., Satterthwaite, T.D., Fan, Y., Launer, L.J., Masters, C.L., Maruff, P., Zhuo, C., Völzke, H., Johnson, S.C., Fripp, J.,Koutsouleris, N., Wolf, D.H., Gur, R., Gur, R., Morris, J., Albert, M.S.,Grabe, H.J., Resnick, S.M., Bryan, R.N., Wolk, D.A., Shinohara, R.T., Shou, H., Davatzikos, C., 2020. Harmonization of large MRI datasets for the analysis of brain imaging patterns throughout the lifespan. NeuroImage 208, 116450. doi:10.1016/j.neuroimage.2019.116450.

Quiñonero-Candela, J., Rasmussen, C.E., 2005. A unifying view of sparse approximate Gaussian process regression. Journal of Machine Learning Research 6, 1939–1959.

Rasmussen, C.E., Williams, C.K.I., 2006. Gaussian Processes for Machine Learning. MIT Press.

Rigby, R.A., Stasinopoulos, D.M., 2004. Smooth centile curves for skew and kurtotic data modelled using the Box–Cox power exponential distribution. Statistics in Medicine 23, 3053–3076. doi:10.1002/sim.1861.

Rigby, R.A., Stasinopoulos, D.M., 2005. Generalized additive models for location, scale and shape. Journal of the Royal Statistical Society: Series C (Applied Statistics) 54, 507–554. doi:10.1111/j.1467-9876.2005.00510.x.

Rousseeuw, P.J., 1984. Least median of squares regression. Journal of the American Statistical Association 79, 871–880. doi:10.1080/01621459.1984.10477105.

Rousseeuw, P.J., Leroy, A.M., 1987. Robust Regression and Outlier Detection. John Wiley & Sons. doi:10.1002/0471725382.

Rousseeuw, P.J., Van Driessen, K., 1999. A fast algorithm for the minimum covariance determinant estimator. Technometrics 41, 212–223. doi:10.1080/00401706.1999.10485670.

Rousseeuw, P.J., van Zomeren, B.C., 1990. Unmasking multivariate outliers and leverage points. Journal of the American Statistical Association 85, 633–639. doi:10.1080/01621459.1990.10474920.

Rutherford, S., Fraza, C., Dinga, R., Wolfers, T., Zabihi, M., Berthet, P., Marquand, A.F., 2022a. Charting brain growth and aging at high spatial precision. eLife 11, e72904. doi:10.7554/eLife.72904.

Rutherford, S., Kia, S.M., Wolfers, T., Fraza, C., Zabihi, M., Dinga, R., Berthet, P., Worker, A., Verdi, S., Andrews, D., et al., 2022b. The normative modeling framework for computational psychiatry. Nature Protocols 17, 1711–1734. doi:10.1038/s41596-022-00696-5.

Sanz-Leon, P., Knock, S.A., Woodman, M.M., Domide, L., Mersmann, J., McIntosh, A.R., Jirsa, V., 2013. The virtual brain: a simulator of primate brain network dynamics. Frontiers in Neuroinformatics 7, 10. doi:10.3389/fninf.2013.00010.

Schwarz, C.G., Gunter, J.L., Ward, C.P., Vemuri, P., Senjem, M.L., Wiste, H.J., Petersen, R.C., Knopman, D.S., Jack, Jr., C.R., 2019. Identification of anonymous MRI research participants with face-recognition software. New England Journal of Medicine 381, 1684–1686. doi:10.1056/NEJMc1908881.

Snoek, J., Larochelle, H., Adams, R.P., 2012. Practical Bayesian optimization of machine learning algorithms, in: Advances in Neural Information Processing Systems.

Stasinopoulos, D.M., Rigby, R.A., 2007. Generalized additive models for location scale and shape (GAMLSS) in R. Journal of Statistical Software 23, 1–46. doi:10.18637/jss.v023.i07.

Stasinopoulos, M.D., Rigby, R.A., Heller, G.Z., Voudouris, V., De Bastiani, F., 2017. Flexible Regression and Smoothing: Using GAMLSS in R. Chapman and Hall/CRC. doi:10.1201/b21973.

Thompson, P.M., Stein, J.L., Medland, S.E., Hibar, D.P., Vasquez, A.A., Renteria, M.E., Toro, R., Jahanshad, N., Schumann, G., Franke, B., et al., 2014. The ENIGMA consortium: large-scale collaborative analyses of neuroimaging and genetic data. Brain Imaging and Behavior 8, 153–182. doi:10.1007/s11682-013-9269-5.

Titsias, M.K., 2009. Variational learning of inducing variables in sparse Gaussian processes, in: Proceedings of the 12th International Conference on Artificial Intelligence and Statistics, pp. 567–574.

Wager, S., Hastie, T., Efron, B., 2014. Confidence intervals for random forests: the jackknife and the infinitesimal jackknife. Journal of Machine Learning Research 15, 1625–1651.

Wang, Y., Li, M., García Reyes, R., Bringas-Vega, M.L., Minati, L., Breakspear, M., Valdés-Sosa, P.A., 2026. The influence of nonlinear resonance on human cortical oscillations. Communications Biology In press.

Wang, Y., Li, M., Paz-Linares, D., Bringas-Vega, M.L., Valdés-Sosa, P.A., 2022. FKreg: A MATLAB toolbox for fast multivariate kernel regression. arXiv preprint arXiv:2204.07716.

Wolfers, T., Doan, N.T., Kaufmann, T., Alnæs, D., Moberget, T., Agartz, I., Buitelaar, J.K., Ueland, T., Melle, I., Franke, B., Andreassen, O.A., Beckmann, C.F., Westlye, L.T., Marquand, A.F., 2018. Mapping the heterogeneous phenotype of schizophrenia and bipolar disorder using normative models. JAMA Psychiatry 75, 1146–1155. doi:10.1001/jamapsychiatry.2018.2467.

Yohai, V.J., 1987. High breakdown-point and high efficiency robust estimates for regression. Annals of Statistics 15, 642–656. doi:10.1214/aos/1176350366.

Yu, M., Linn, K.A., Cook, P.A., Phillips, M.L., McInnis, M., Fava, M., Trivedi, M.H., Weissman, M.M., Shinohara, R.T., Sheline, Y.I., 2018. Statistical harmonization corrects site effects in functional connectivity mega-analysis. Human Brain Mapping 39, 4213–4227. doi:10.1002/hbm.24241.

Zabihi, M., Oldehinkel, M., Wolfers, T., Frouin, V., Goyard, D., Loth, E., Charman, T., Tillmann, J., Banaschewski, T., Dumas, G., Holt, R.J., Baron-Cohen, S., Durston, S., Bölte, S., Murphy, D., Marquand, A.F., Buitelaar, J.K., 2019. Dissecting the heterogeneous cortical anatomy of autism spectrum disorder using normative models. Biological Psychiatry: Cognitive Neuroscience and Neuroimaging 4, 567–578. doi:10.1016/j.bpsc.2018.11.013.

Zhu, L., Liu, Z., Han, S., 2019. Deep leakage from gradients, in: Advances in Neural Information Processing Systems.

